# Thioesterase modulates antibacterial Type VI secretion system via alteration in oxidative stress responses in *Burkholderia*

**DOI:** 10.64898/2026.10.05.756662

**Authors:** Rajni Yadav, Kristi Kabyashree, Rahul Kumar, Mandira Roy, Gopaljee Jha

## Abstract

Bacteria utilize specialized protein secretion systems, particularly the type VI secretion system (T6SS), to dominate interbacterial competitions and kill the prey bacteria in a contact-dependent manner. Here, we report that disruption of the *thioesterase* gene encoded in the gramibactin (a bacterial siderophore) biosynthetic gene cluster activates the T6SS in the *Burkholderia gladioli* strain NGJ1, even in the absence of prey. The *thioesterase* mutant (*Δthio*) exhibits enhanced antibacterial activity, and ectopic expression of the gene complements the phenotype. Notably, *Δthio* mutant as well as *thioesterase*-complement strains show enhanced sensitivity to oxidative stress. The qRT-PCR analysis showed that the *Δthio* mutant had upregulated expression of RpoN1, a sigma factor, which positively regulates T6SS functions. We demonstrate that *ΔrpoN1* is hypersensitive to oxidative stress and defective in T6SS activation, whereas overexpression of RpoN1 enhances oxidative stress tolerance in NGJ1 and activates T6SS. We propose that, during interbacterial conflicts encountered during plant colonization, NGJ1 faces an oxidative-stress-enriched environment that modulates *thioesterase* expression, induces *RpoN1*, and activates T6SS. Overall, our study exemplifies the broader physiological role of Thioesterase in maintaining intracellular ROS homeostasis in bacteria and stimulating antibacterial activity.

## Introduction

Type VI secretion system (T6SS) is known to regulate several physiological processes in bacteria, including commensalism or symbiosis (Serapio-Palacios et al., 2022; Speare et al., 2018) biofilm formation (Gallique et al., 2017; Teschler et al., 2022; Yang et al., 2022), oxidative stress responses (Li et al., 2024a; Si et al., 2017; Zuo et al., 2023), quorum sensing (Alves et al., 2022; Kumar et al., 2025; Lesic et al., 2009; Mashruwala et al., 2022; Shao et al., 2018), iron acquisition (Haas et al., 2023; Lin et al., 2017a, 2017b), and osmotic stress (Yang et al., 2022; Yin et al., 2024). The regulation of T6SS is complex and occurs at transcriptional, post-transcriptional, and post-translational levels. At the transcriptional level, alternative sigma factor RpoN (σ□□) plays a central role in regulating T6SS expression in several bacteria, including *Vibrio cholerae*, *Pseudomonas aeruginosa*, and *Klebsiella pneumoniae* (Allsopp et al., 2022; Basler et al., 2013; Dong & Mekalanos, 2012; Gu et al., 2022; Hendrickson et al., 2001; Shao et al., 2018; Storey et al., 2020). Post-transcriptionally, the GacS/ GacA two-component system modulates small regulatory RNAs (such as RsmX, RsmY, and RsmZ), which sequester RNA-binding proteins, such as RsmA/ RsmE, that repress translation of T6SS components (Allsopp et al., 2017; Dupont et al., 2025; Lapouge et al., 2008). At the post-translational level, T6SS is regulated by the threonine phosphorylation pathway (TPP) and other independent pathways in *P. aeruginosa* (George et al., 2024; Hsu et al., 2009; Mougous et al., 2007). Evidence suggests that microbial secondary metabolites can influence antibacterial activity. These metabolites are typically encoded by biosynthetic gene clusters (BGCs), which can now be systematically identified using genome mining tools such as antiSMASH and PRISM (Blin et al., 2017; Skinnider et al., 2020). While not essential for primary growth, secondary metabolites play crucial roles in microbial defense, signalling, and ecological adaptation (Tyc et al., 2017). For instance, glutathione sensing via the VirA/ VirG system activates the T6SS in *Burkholderia pseudomallei* (Ku & Gan, 2019; Wong et al., 2015), whereas pyrroloquinoline quinone (PQQ) modulates T6SS activity in *Serratia marcescens* through pH changes (Jia et al., 2024). Similarly, T6SS has been linked to the regulation or secretion of phenazine compounds in *P. chlororaphis* (Liu et al., 2018). Despite these insights, the mechanistic interplay between secondary metabolites and T6SS regulation remains largely unexplored.

Previously, we have reported that an endophytic bacterium, *Burkholderia gladioli* NGJ1, isolated from rice, demonstrates strong antifungal as well as mycophagous (fungal eating) (Swain et al., 2017) and antibacterial activities (Yadav et al., 2021). It has been demonstrated that functional T6SS is required for contact-dependent antibacterial activity, while the T3SS (Type III secretion system) is required for mycophagous ability. In this study, we attempted to identify and characterize the involvement of secondary metabolite-encoding BGCs with respect to the antifungal and antibacterial properties of NGJ1. We report that disruption of the *thioesterase* gene, encoded in the gramibactin biosynthetic cluster, compromises the mycophagous ability of NGJ1. On the other hand, it leads to constitutive activation of T6SS, thereby enhancing the antibacterial activity of NGJ1. We observed that this shift is associated with upregulation of the sigma factor (*rpoN1*), T6SS-associated genes and effectors, and oxidative stress-responsive genes. We propose that downregulation of *thioesterase* induces oxidative stress in bacteria, triggering the *rpoN1*-dependent regulatory network that promotes T6SS activation. Overall, our study presents a functional link between Thioesterase and T6SS regulation, and exemplifies how metabolic networks shape microbial interactions.

## Results

### BGCs are important for the mycophagous ability of the *Burkholderia gladioli* strain NGJ1

Using the bioinformatic tool (antiSMASH 5), we identified thirteen Biosynthetic Gene Clusters (BGCs) that encode secondary metabolites in NGJ1. Amongst them, Isobongkrekic acid, Enacyloxin IIa, Orfamide B, Gramibactin, and Rhizomide A/B/C were selected to explore their involvement in mycophagy (Supplementary Table S1; Supplementary Fig S1). The expression of a few genes from each of these BGCs [Isobongkrekic acid (*Polyketide beta-ketoacyl ACP synthase, Pbk-ACPs*) and *Cytochrome P450* (*CYP450*), Enacyloxin IIa: *Mate family efflux transporter* (*MFET*) and *LuxR family Transcription regulator* (*LuxR*), Orfamide A: *Hemolysin secretion protein D (HlyD)*, Gramibactin [*Thioesterase* (*Thio*)*, MFS transporter* (*MFS*)] and Rhizomide A/B/C: *Acyl carrier protein* (*ACP*) was analyzed using qRT-PCR and data revealed that they were upregulated during 24h of interaction with *R. solani* (Supplementary Fig S2), compared to the expression during NGJ1 growth in PDB (with *R. solani*). Further, using *pCZ367:*:*lac*Z plasmid integration, we attempted to create *lacZ* insertion mutations (GUS-tagged) in these genes and obtained insertions in *Pbk-ACPs*, *CYP450*, *MFET*, *LuxR*, *Thio,* and *MFS* genes. We observed that each of these mutants was GUS-positive and compromised in its ability to prevent *R. solani* growth on PDA plates (Supplementary Fig. S3A). On the other hand, as reported previously (Swain et al. 2017), wild-type (WT) NGJ1 completely prevented the fungal growth. Only a few *R. solani* sclerotia were produced on PDA plates during confrontation with WT, whereas a large number of secondary sclerotia (in hundreds) were formed during confrontation with the mutant strains (Supplementary Fig. S3B). Also, most of the secondary sclerotia were viable when grown on fresh plates (Supplementary Fig. S3C).

Further, the antibacterial activity of the mutants was tested against *Escherichia coli* and *Xanthomonas oryzae* pv. oryzae (the causal agent of bacterial leaf blight disease in rice). It was intriguing that the zone of inhibition was higher in mutant strains than in WT (Supplementary Fig. S4). As the mutants were created by plasmid integration, the observed phenotype may be due to polar effects on downstream genes and to overall defects in the synthesis of secondary metabolites from the BGCs. To investigate further, we selected Thioesterase (a key catalytic enzyme involved in the processing and off-loading step of Gramibactin from the NRPS assembly) and the MFS transporter (a Major Facilitator Superfamily transporter that mediates the export of the synthesized Gramibactin to the extracellular milieu) to examine the underlying mechanism of enhanced antibacterial activity.

### Loss of Thioesterase leads to constitutive activation of T6SS-mediated antibacterial activity in NGJ1

NGJ1 is known to kill prey bacteria in a contact-dependent manner (Yadav et al., 2021). We observed that compared to WT, the Δ*thio* mutant (*thio::*lacZ) exhibited significantly enhanced antibacterial activity when co-cultured with prey (*E. coli*) bacteria (Fig. 1A). On the other hand, antibacterial activity of the Δ*mfs* mutant (*mfs::*lacZ) was comparable to that of NGJ1 (Fig. 1A). Notably, the ectopic expression of the full-length *thioesterase* gene restored the antibacterial activity of the complement strain (Δ*thio +* pHM1:*thio*) to WT level. The *thioesterase*-complement strain was also proficient in mycophagy (Supplementary Fig. S5A) and in preventing sheath blight disease, caused by *R. solani* in rice (Supplementary Fig. S5B and S5C), while the Δ*thio* mutant was compromised in these processes. Notably, Δ*thio* as well as *thioesterase*-complement strain were defective in siderophore production (Supplementary Fig. S6). This suggested that the Gramibactin BGC is involved in siderophore production, and the siderophore-deficient phenotype of the Δ*thio* strain is due to polar effects on downstream genes in the biosynthetic pathway. On the other hand, enhanced contact-dependent antibacterial activity of Δ*thio* mutant is predominantly due to defects in Thioesterase function and not due to the polar effects on Gramibactin biosynthesis, as it was fully complemented by expression of *thioesterase* gene.

**Figure 1.**
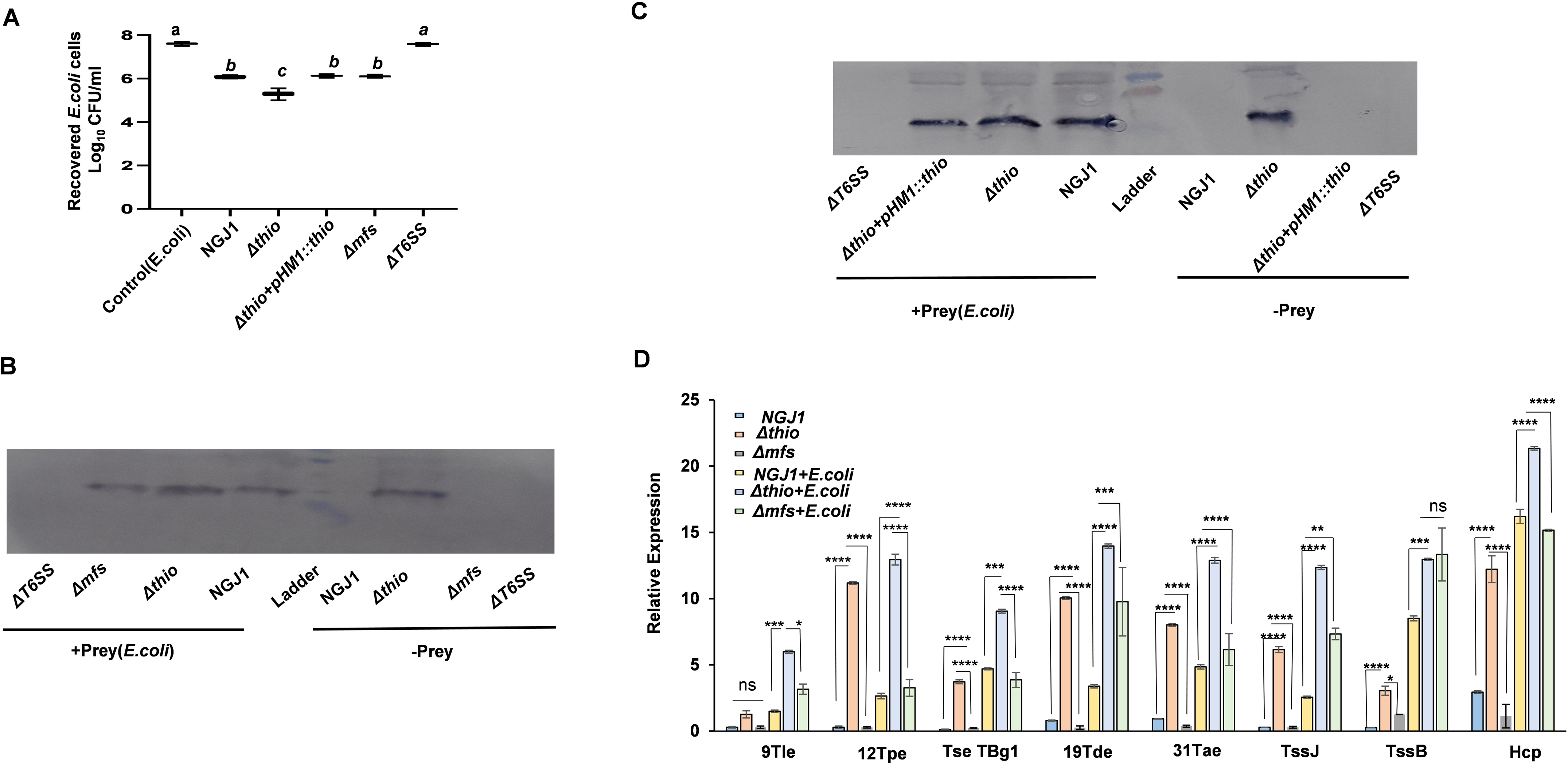
Disruption of *thioesterase* enhances T6SS-mediated antibacterial activity of *B. gladioli* strain NGJ1 against *E. coli*. **A)** Contact-dependent antibacterial activity of NGJ1 against prey (*E. coli*), estimated as colony-forming units (CFU) of the recovered prey bacterial cells. The whiskers depict the minimum and maximum values. Different letters indicate significant difference (one-way ANOVA, Tukey’s multiple comparisons post hoc test; p<0.05). **B)** and **C)** Immunoblot showing secretion of Hcp-1 protein (associated with T6SS-1 of NGJ1) by different NGJ1 strains, when grown in PDB, with and without prey bacteria, using protein-specific antibody. **D)**The qRT-PCR-based expression of genes encoding structural components (TssJ, TssB, Hcp-1) and effectors (9Tle, 12Tpe, TseTBg1, 19Tde, and 31Tae) of T6SS during 24h of NGJ1 growth in the presence/ absence of prey, using the 16S rRNA gene as endogenous control. Data was estimated using (two-way ANOVA, Tukey’s multiple comparisons test) and Asterisks ^∗, ∗∗, ∗∗∗, ∗∗∗∗^ indicate a significant difference at p<0.5, p<0.05, p<0.001, and p<0.0001, respectively. ‘ns’ indicates a non-significant difference. The experiments were independently repeated three times, with a minimum of 3 technical replicates. Graphs show mean values ± standard error of three biological replicates.

Considering that the Type VI secretion system (T6SS) is involved in contact-dependent antibacterial activity of NGJ1 (Yadav et al., 2021), as a hallmark for T6SS activation, we analyzed Hcp-1 (associated with T6SS-1 of NGJ1) secretion by different NGJ1 strains using immunoblotting with the specific antibody, as reported previously (Yadav et al., 2021). Notably, in the absence of the prey bacteria (*E. coli*), Δ*thio* mutant, but not the Δ*mfs/* WT strains, secreted the Hcp proteins into the extracellular milieu (Fig. 1B; Supplementary File 1). The *thioesterase*-complement strain behaved like WT and failed to secrete Hcp-1 in the absence of the prey (Fig. 1C; Supplementary File 2). On the other hand, WT, Δ*thio*, *thioesterase*-complement as well as Δ*mfs* strains were capable of secreting Hcp-1, only in the presence of the prey (Fig. 1B). The previously reported T6SS mutant of NGJ1 was used as a negative control in immunoblot analysis. Overall, these analyses emphasize that disruption of Thioesterase function leads to constitutive activation of the T6SS and that the phenotype is complemented by ectopic expression of the gene.

To further elucidate, we analyzed the expression of the T6SS-1 apparatus (TssJ, TssB, Hcp) and effectors (9Tle, 12Tpe, TseTBg1, 19Tde, 31Tae) encoding genes (Yadav et al., 2021) in different NGJ1 strains in the presence or absence of the prey at 24h. The qRT-PCR analysis showed that most of the tested genes had a basal level of expression in WT/ D*mfs* mutant and were upregulated in the D*thio* mutant when grown in the absence of the prey (Fig. 1D). In the presence of the prey, the upregulation of these genes was further enhanced in D*thio* mutant (Fig. 1D). Taken together, our data indicate that the D*thio* mutant had a constitutively active T6SS (even in the absence of prey), which gets further stimulated in the presence of prey.

It was noteworthy that the Δ*thio* mutant also exhibited enhanced contact-dependent antibacterial activity against *X. oryzae* pv. oryzae, while the antibacterial activity of the complement strain (Δ*thio+*pHM1::*thio*) was comparable to that of WT (Fig. 2A). Moreover, compared with the WT and complement strains, the Δ*thio* mutant led to a significant reduction in the disease severity, as reflected in reduced lesion length caused by *X. oryzae* pv. oryzae in rice (Fig. 2B and 2C).

**Figure 2.**
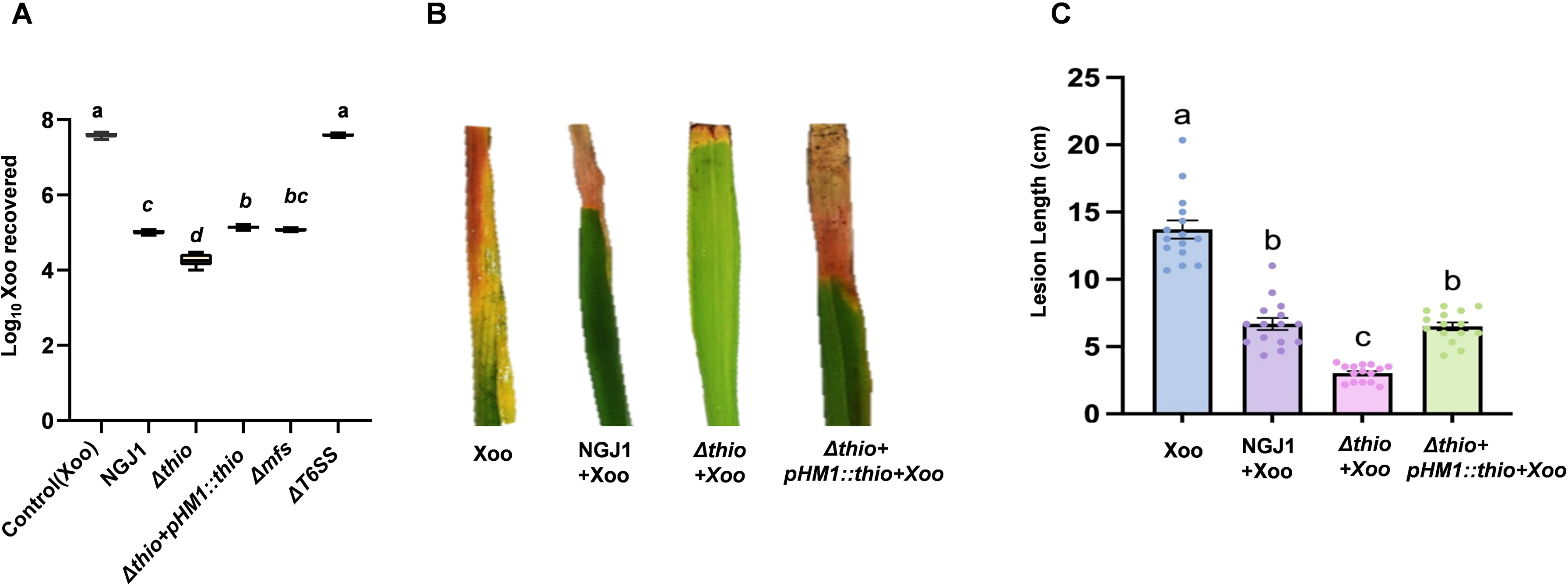
Disruption of *thioesterase* enhances antibacterial activity of *B. gladioli* strain NGJ1 against the bacterial leaf blight pathogen, *Xanthomonas oryzae* pv. oryzae. A) Contact-dependent antibacterial activity of NGJ1 against *X. oryzae* pv. oryzae (Xoo, prey), estimated as colony-forming units (CFU) of the recovered prey bacterial cells. The whiskers depict the minimum and maximum values; Different letters indicate significant difference (one-way ANOVA, Tukey’s multiple comparisons post hoc test; p<0.05). **B)** Representative images of bacterial leaf blight symptoms in rice elicited by *Xoo* (control), and *Xoo* along with NGJ1 strains. **C)** Graph showing lesion lengths measured at 14 days of post-inoculation. Data were analyzed using one-way ANOVA and Tukey’s multiple comparisons post hoc test; p<0.05. The experiments were independently repeated three times, with 15 plants analyzed per treatment. Graphs show mean ± standard error of three biological replicates. Dot represents individual data points.

### Disruption of Thioesterase alters the expression of global transcriptional regulators in NGJ1

Thioesterase hydrolyzes the thioester bond between a carbonyl group and a sulfur. It catalyzes reactions in pathways such as fatty acid, polyketide, and non-ribosomal peptide synthesis (Cantu et al., 2010). The available literature is unclear about how the Thioesterase can modulate T6SS functions in bacteria. Therefore, to get further insights into T6SS activation in the D*thio* mutant, we examined the expression of global master transcription regulators, including GacA (global activator in GacA/GacS two-component regulatory system (Tejada-Arranz et al., 2025), RsmA (associated with post-transcriptional repression of T6SS (Allsopp et al., 2017), RpoN (Allsopp et al., 2022b, Das et al.,2022), and OxyR (Li et al., 2024b; Si et al., 2017). The qRT-PCR analysis reflected that during growth in PDB, D*thio* mutant had upregulated expression of *oxyR* and *rpoN1*, and downregulated expression of *gacA*, *rsmA,* and *rpoN2* genes, compared to WT (Fig. 3A). The *thioesterase*-complement strain had downregulated expression of *rpoN1* and upregulated expression of *gacA*, *rsmA,* and *rpoN2* genes, similar to WT. Notably, both D*thio* and *thioesterase*-complement strains had enhanced *oxyR* expression, compared to WT. Taken together, our data indicated that disruption of *thioesterase* activates the RpoN1/OxyR regulatory pathway and represses the GacA/RsmA pathway in NGJ1.

**Figure 3.**
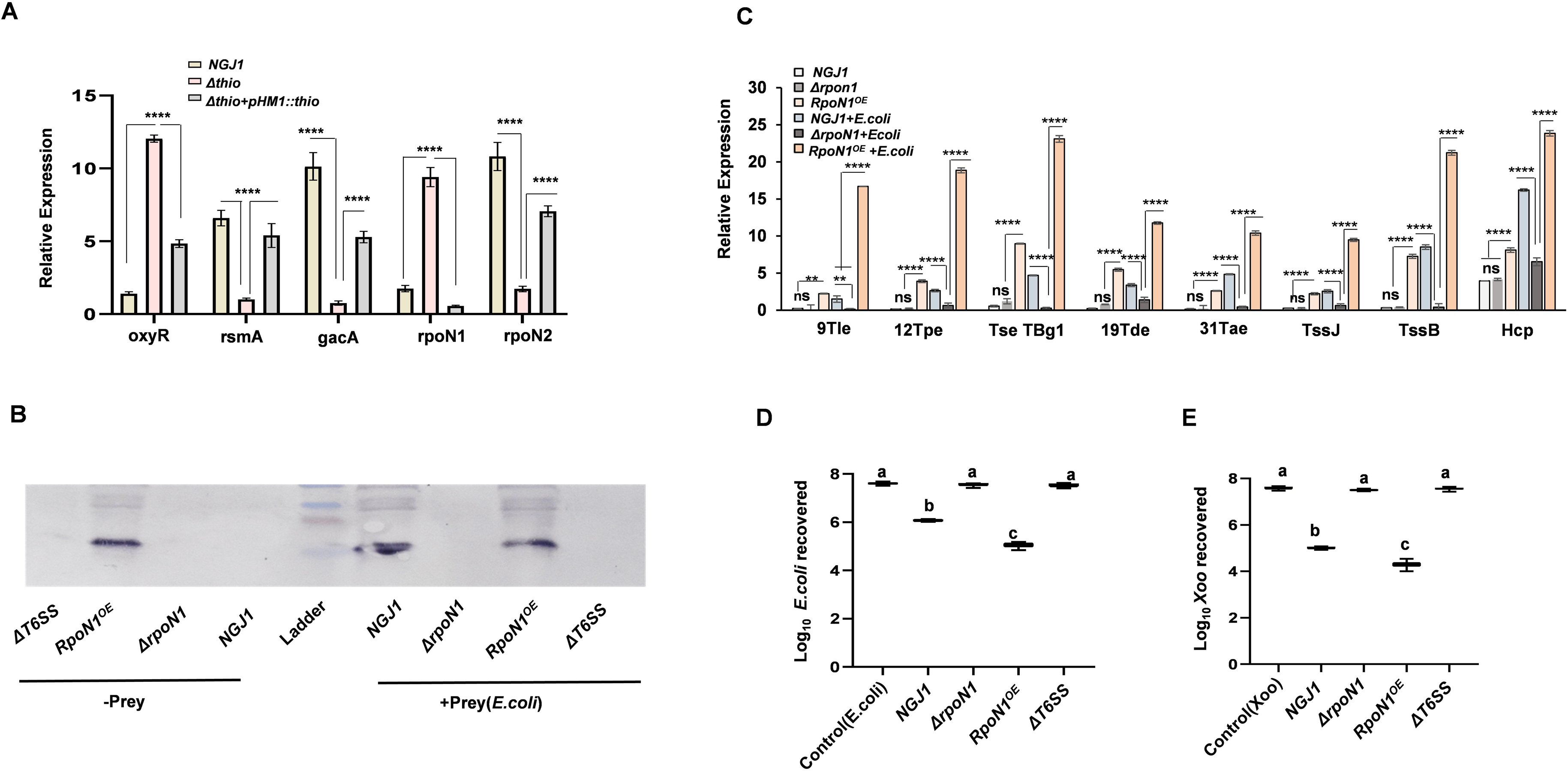
Thioesterase modulates T6SS-mediated antibacterial activity by repressing *rpoN1* expression. **A)** qRT-PCR-based expression of *oxyR*, *rsmA*, *gacA*, *rpoN1*, and *rpoN2* genes in different strains of NGJ1, at 24h of growth. The relative expression was estimated using the 16S rRNA of NGJ1 as an endogenous control. Asterisks ^∗, ∗∗,∗∗∗, ∗∗∗∗^ indicate a significant difference at p<0.5, p<0.05, p<0.001, and p<0.0001, respectively (estimated using two-way ANOVA, Tukey’s multiple comparisons test). **B)** Immunoblot using protein-specific antibody showing secretion of Hcp-1 by different strains of NGJ1, when grown in the presence/ absence of the prey bacteria (*E. coli*). **C)** qRT-PCR-based expression of T6SS apparatus (TssJ, TssB, Hcp), and effector (9Tle, 12Tpe, TseTBg1, 19Tde, and 31Tae) encoding genes of NGJ1 at 24h of growth in the presence and absence of prey bacteria. Relative expression was estimated using 16S rRNA from NGJ1 as an endogenous control. Asterisks ^∗, ∗∗, ∗∗∗, ∗∗∗∗^ indicate a significant difference at p<0.5, p<0.05, p<0.001, and p<0.0001, respectively (estimated using two-way ANOVA, Tukey’s multiple comparisons test). ‘ns’ indicates a non-significant difference between indicated samples. **D)** and **E)** contact-dependent antibacterial activity of NGJ1 strains against prey bacteria (**D**. *E. coli* and **E**. *Xoo*), estimated as colony-forming units (CFU) of the recovered prey bacterial cells. The whiskers depict the minimum and maximum values; Different letters indicate significant difference (one-way ANOVA, Tukey’s multiple comparisons post hoc test; p<0.05). The experiments were independently repeated three times, with a minimum of 3 technical replicates per condition. The graph shows the mean ± standard error of three biological replicates.

### Thioesterase regulates T6SS functions via modulation of RpoN1

We used the previously reported *rpoN1* mutant (D*rpoN1*) of NGJ1 (Das et al., 2022) and generated an *rpoN1* overexpression (NGJ1 + pHM1::*RpoN1, RpoN1^OE^*) strain to understand its role in T6SS activation. The immunoblot analysis reflected that *RpoN1^OE^* strain translocated Hcp-1 into the extracellular milieu, even in the absence of the prey. The D*rpoN1* mutant was unable to secrete Hcp-1, whereas WT secreted it only when grown in the presence of the prey (Fig. 3B; Supplementary File 3). This suggested that RpoN1 is important for the T6SS activation in NGJ1. The qRT-PCR analysis also reflected that the T6SS-1 apparatus and effector-encoding genes were expressed at a basal level in WT and mutant strains, whereas they were upregulated in RpoN1^OE^, when grown in the absence of prey (Fig. 3C). In the presence of prey, upregulation of the T6SS-1 apparatus and T6SS effectors was significantly higher in RpoN1^OE^ compared to WT (Fig. 3C). Moreover, we observed that RpoN1^OE^ was proficient, whereas the Δ*rpoN1* and Δ*T6SS* strains were defective in contact-dependent antibacterial activity against *E. coli* (Fig. 3D) *and X. oryzae* pv. oryzae (Fig. 3E). These results indicated that RpoN1 is a positive regulator of T6SS-mediated antibacterial activity in NGJ1.

### Thioesterase modulates oxidative stress response

The elevated *oxyR* expression in the D*thio* mutant and complement strains prompted us to investigate whether Thioesterase is involved in oxidative stress management in NGJ1. We tested the sensitivity of different NGJ1 strains against 5- and 20-mM H₂O₂ treatment on PDA plates. Notably, D*thio, thioesterase*-complement, and D*rpoN1* strains exhibited enhanced sensitivity to 20 mM H₂O₂, with significantly larger zones of inhibition than those for WT and RpoN1^OE^ (Fig. 4A and Supplementary Fig. S7A). The confocal microscopy analysis of H_2_DCFDA (Dichlorodihydrofluorescein diacetate, a ROS-sensitive dye) stained bacterial cells showed that fluorescence signal (reflecting ROS accumulation) was not detected when grown in PDB (Supplementary Fig. S7B). Upon H₂O₂ treatment, only a limited number of WT and RpoN1^OE^ cells showed signal, whereas a large number of fluorescent cells were observed in D*thio*, D*rpoN1*, and *thioesterase*-complement strains (Fig. 4B, Supplementary Fig. S7C). Further, we quantified intracellular ROS accumulation by measuring dichlorofluorescein fluorescence intensity signal using a spectrophotometer, and observed that under oxidative stress (H₂O₂ treatment), compared to WT and RpoN1^OE^ strains, D*thio*, D*rpoN1*, and *thioesterase*-complement strains had significantly higher ROS levels (Fig. 4C). Moreover, compared to WT and RpoN1^OE^, *oxyR* expression was significantly higher in Δ*thio*, *thioesterase*-complement, and Δ*rpoN1* strains (Fig. 4D). Overall, these analyses suggested that D*rpoN1* and D*thio* mutants, as well as *thioesterase*-complement strains, have enhanced sensitivity to oxidative stress.

**Figure 4.**
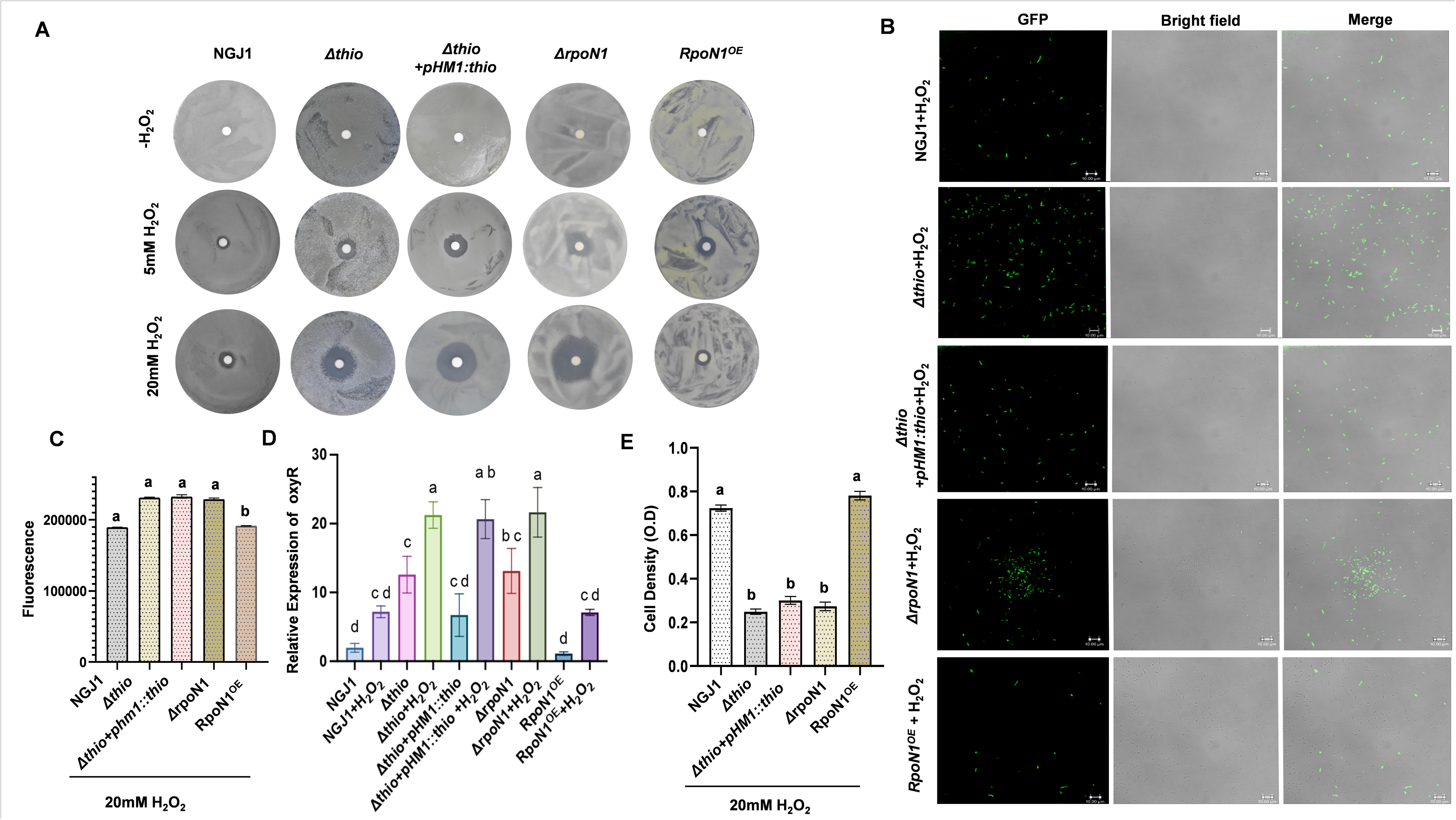
Thioesterase modulates OxyR-dependent oxidative stress responses in *B. gladioli* NGJ1. **A)** Sensitivity of different NGJ1 strains against 5- and 20mM H_2_O_2_ when grown for 48h on PDA plates. **B)** Confocal micrographs of DCFDA-stained NGJ1 strains after 24h of 20mM H_2_O_2_-treatment. Fluorescent cells reflect the accumulation of reactive oxygen species (ROS). Scale bars = 10 µm. **C)** Intracellular ROS level in 20mM H_2_O_2_-treated DCFDA-stained NGJ1 strains, quantified by a spectrophotometer (excitation: 485nm and emission: 535nm). Statistical significance was analyzed using one-way ANOVA and Tukey’s multiple-comparisons post hoc test (p<0.05). Different letters indicate a significant difference. **D)** qRT-PCR-based expression of *oxyR* in different strains of NGJ1, during growth in PDB, with and without H_2_O_2_ treatment. Relative expression was estimated using the 16S rRNA of NGJ1 as an endogenous control. Data were analyzed using one-way ANOVA and Tukey’s multiple comparisons post hoc test. Asterisks ^∗, ∗∗, ∗∗∗, ∗∗∗∗^ indicate a significant difference at p<0.5, p<0.05, p<0.001, and p<0.0001, respectively. ‘ns’ indicates a non-significant difference between the indicated groups. **E)** The concentration of NGJ1 cells after 24h of 20mM H_2_O_2_ treatment, estimated as OD^595nm^. Data were analyzed using one-way ANOVA and Tukey’s multiple comparisons post hoc test (p<0.05). Different letters indicate a significant difference. The graph shows the mean ± standard error across three biological replicates. The experiments were independently repeated three times, with a minimum of 3 technical replicates per condition.

We also quantified the bacterial cell concentrations by measuring the absorbance (O.D) at 595nm and observed that compared to WT and RpoN1^OE^, the population of Δ*thio*, complement strain, and Δ*rpoN1* strains was reduced when grown in the presence of 20 mM H₂O₂ (Fig. 4E). The D*mfs* mutant had marginal sensitivity to H₂O₂ (Supplementary Fig. S8C), and its growth was comparable to WT under oxidative stress (Supplementary Fig. 8D).

To further investigate the role of Thioesterase in oxidative stress management, we ectopically overexpressed it in *E. coli* (BL-21) using an IPTG-inducible pET28a-based bacterial system. Upon IPTG induction, the strain’s sensitivity to 20 mM H₂O₂ was assessed, and it was observed that overexpression of *thioesterase* (*thioesterase*^OE^) confers enhanced sensitivity to oxidative stress, as OD^595^ (Fig. 5A), and colony-forming unit, CFU (Fig. 5B) were significantly reduced in H₂O₂-treated *thioesterase*^OE^ strain, as compared to the control strain with empty vector (pET28a). Moreover, compared with the control, a larger proportion of *thioesterase*^OE^ cells showed greater ROS accumulation (visualized as green fluorescence by H_2_DCFDA staining and confocal microscopy) upon H₂O₂ treatment (Fig. 5C and 5D).

**Figure 5.**
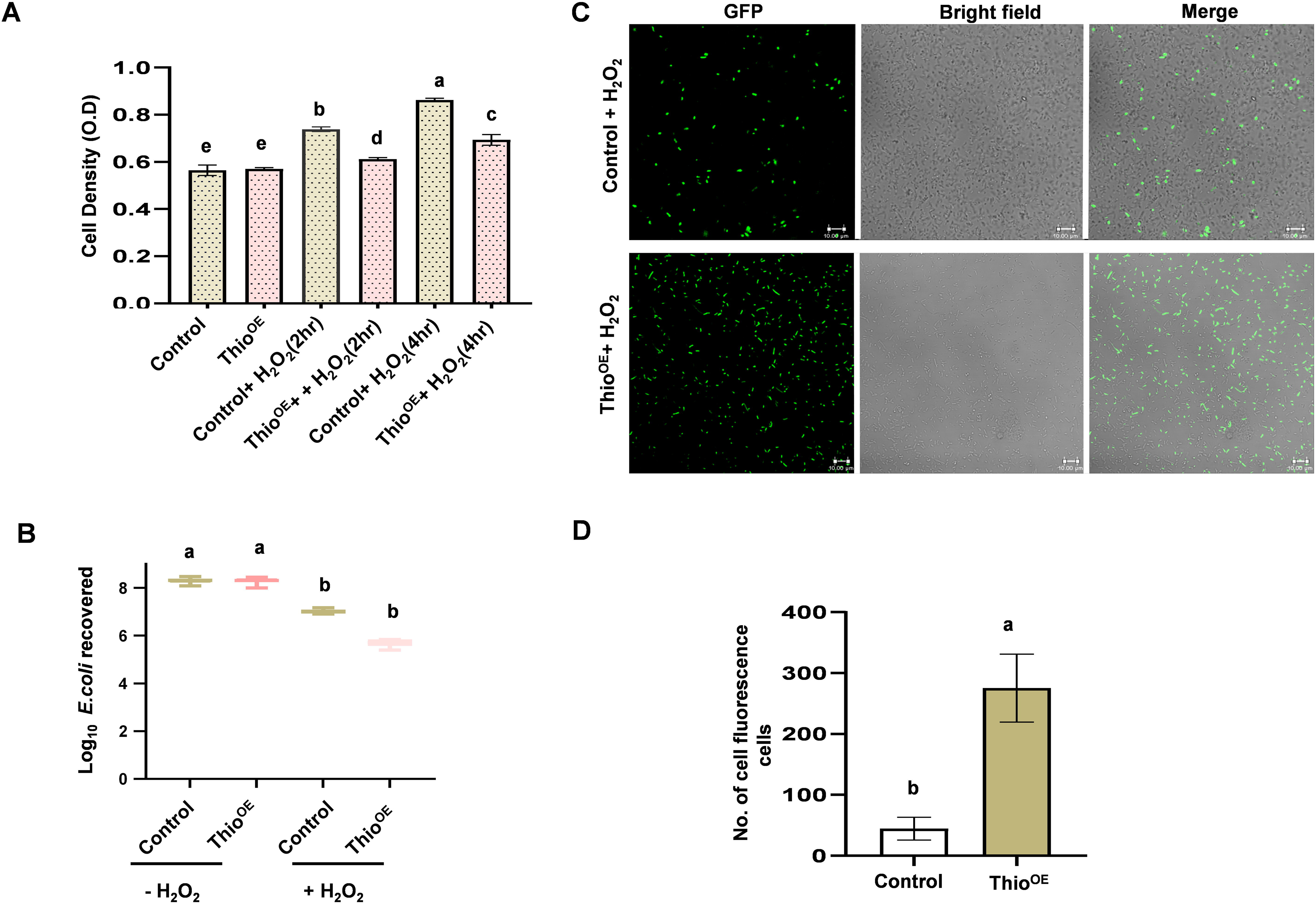
Heterologous expression of the *thioesterase* gene of NGJ1 provides oxidative-stress tolerance in *E. coli*. **A)** The concentration of *E. coli* cells expressing the *thioesterase* gene of NGJ1 (Thio^OE^) and vector control (pET-28a), estimated as OD^595nm^, upon 2 and 4h of 20mM H_2_O_2_ treatment. **B)** The colony-forming units (CFU) of the *E. coli* cells, recovered with and without 20mM H_2_O_2_ treatment. The whiskers depict the minimum and maximum values. Different letters indicate significant difference (one-way ANOVA, Tukey’s multiple comparisons post hoc test; p<0.05). **C)** The confocal micrographs showing ROS accumulation (fluorescing cells) in DCFDA-stained 20mM H_2_O_2_-treated *E. coli* cells. Scale bars = 10 µm. **D)** Graph showing the number of fluorescent *E. coli* cells upon 20mM H_2_O_2_ treatment. Similar results were obtained in at least 3 independent replicates, and only representative images are shown. Data were analyzed using one-way ANOVA and Tukey’s multiple comparisons post hoc test (p<0.05); and different letters indicate significant difference (p<0.05). The graph shows the mean ± standard error of three biological replicates. The experiments were independently repeated three times, with a minimum of 3 technical replicates per condition.

Overall, our analysis suggested that both overexpression and disruption of *thioesterase* enhance ROS accumulation in the bacteria, rendering them hypersensitive to oxidative stress.

### Thioesterase-mediated oxidative stress environment modulates T6SS activation

We further envisaged studying the physiological relevance of Thioesterase-mediated sensitivity to oxidative stress, with respect to contact-dependent antibacterial activity and T6SS activation in NGJ1. To test this, we subjected different strains of NGJ1 to oxidative stress (20 mM H_2_O_2_) and observed that under oxidative stress conditions, the WT, D*thio*, and *thioesterase*-complement NGJ1 strains translocated Hcp-1 into the extracellular milieu, whereas D*T6SS-1* was defective in this process (Fig. 6A; Supplementary File 4). However, under normal growth conditions (without H_2_O_2_ treatment), only the D*thio* mutant secretes Hcp-1. We also measured *rpoN1* expression and observed that it is induced under oxidative stress across different NGJ1 strains (except Δ*rpoN1*); however, compared with the others, the Δ*thio* and *RpoN1*^OE^ strains showed significantly higher *rpoN1* expression. This suggests that increased *rpoN1* expression leads to T6SS activation. This is consistent with our immunoblot analysis, which showed that Δ*thio* and RpoN1^OE^ strains had autoactivated T6SS, even in the absence of prey bacteria, whereas Δ*rpoN1* is defective in T6SS.

**Figure 6:**
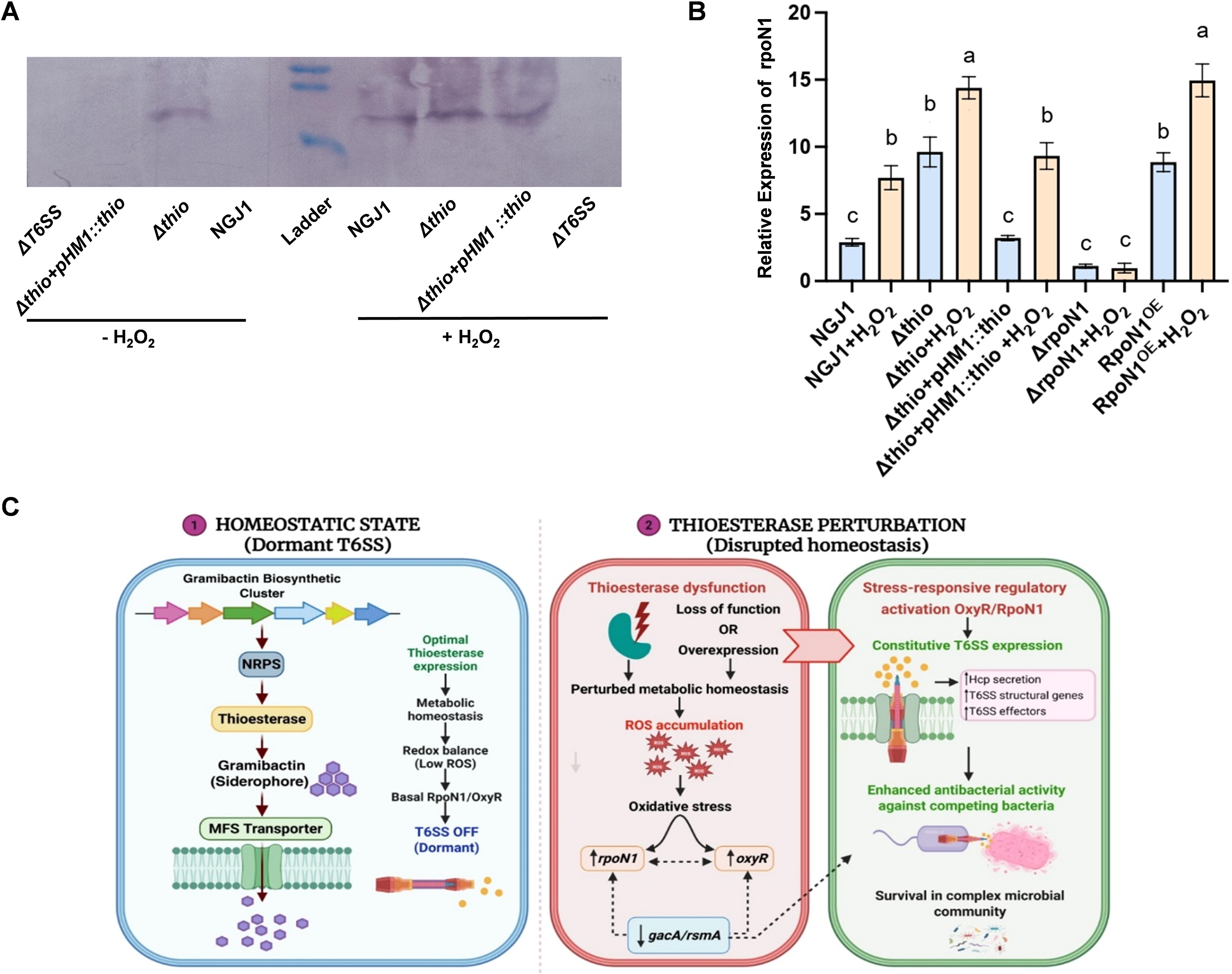
Thioesterase mediates oxidative stress tolerance and constitutive activation of T6SS in *B. gladioli* strain NGJ1. **A)** Immunoblot showing secretion of Hcp-1 (associated with T6SS-1) protein by different NGJ1 strains upon 20mM H_2_O_2_ treatment (24h), using protein-specific antibody. **B** and **C)** qRT-PCR-based expression of *rpoN1* genes in different strains of NGJ1, during growth in PDB, with and without H_2_O_2_ treatment. Relative expression was estimated using the 16S rRNA of NGJ1 as an endogenous control. Data were analyzed using one-way ANOVA and Tukey’s multiple comparisons post hoc test. Asterisks ^∗, ∗∗, ∗∗∗, ∗∗∗∗^ indicate a significant difference at p<0.5, p<0.05, p<0.001, and p<0.0001, respectively. ‘ns’ indicates a non-significant difference between the indicated groups. The experiments were independently repeated three times, with a minimum of 3 technical replicates per condition. **D)** Proposed model for *thioesterase*-mediated modulation of T6SS-dependent antibacterial activity in NGJ1. Thioesterase helps maintain redox balance and prevents T6SS activation under normal growth conditions. Therefore, disruption of Thioesterase function enhances ROS accumulation and creates an oxidative stress-enriched environment, inducing *rpoN1* (sigma transcription factor) and *oxyR* (Oxidative stress regulator) expression. The elevated level of these master regulators results in constitutive activation of the T6SS and enhanced antibacterial activity, highlighting the role of Thioesterase in coordinating oxidative stress adaptation and T6SS regulation in NGJ1.

Overall, we propose that disruption of Thioesterase function enhances ROS accumulation and creates an oxidative stress-enriched environment that induces RpoN1 (sigma transcription factor) expression, leading to constitutive activation of the T6SS and enhanced antibacterial activity (Fig. 6C).

## Discussion

Previous studies have suggested that bacteria deploy various protein secretion systems, particularly the type VI secretion system (T6SS), to dominate inter-bacterial conflicts, acquire nutrients, and adapt to stress during host colonization (Rivard et al., 2026; Wohlfarth et al., 2026). Also, diverse antimicrobial secondary metabolites, siderophores, and other signaling molecules play important roles during bacterial interactions with their hosts and other microbes (Kramer et al., 2020; Wei et al., 2025). The coordination between T6SS and secondary metabolites in modulating bacterial traits largely remains uncharacterized. In this study, we demonstrate that a bacterial siderophore, Gramibactin-associated Thioesterase, acts as a determinant of T6SS-dependent antibacterial activity and regulates redox homeostasis in *Burkholderia gladioli* strain NGJ1.

NGJ1 bacterium is known to activate its T6SS and deploy its antibacterial effectors to dominate interbacterial conflicts (Yadav et al. 2021). However, *thioesterase* mutant (Δ*thio*) NGJ1 had T6SS activated, even in the absence of prey bacteria, whereas disruption of a downstream gene (*MFS transporter*) of its biosynthetic gene cluster (gramibactin BGC) failed to trigger T6SS. Moreover, Δ*thio*, but not the *mfs* mutant (Δ*mfs*), demonstrated enhanced contact-dependent antibacterial activity against *Xanthomonas oryzae* and *E. coli*. Also, the Δ*thio* mutant phenotype was complemented by expression of the full-length gene. Thioesterase of NGJ1 belongs to type II Thioesterases, and it is involved in polyketide biosynthesis (Geyer et al., 2022; Wang et al., 2022). Therefore, we conclude that, in addition to its canonical role in metabolite biosynthesis, Thioesterase modulates T6SS functions.

Notably, both the Δ*thio* mutant and the *thioesterase*-complement strain were more sensitive to oxidative stress than the WT. Also, heterologous expression of *thioesterase* rendered *E. coli* cells sensitive to oxidative stress. This tempted us to speculate that both loss and gain of Thioesterase function may modulate ROS homeostasis in bacteria. Alterations in redox homeostasis led to transcriptional changes in bacteria and modulate T6SS functions (Hespanhol et al., 2023; Reniere, 2018; Sevilla et al., 2019). We observe that the expression of genes encoding OxyR, an oxidative-stress-responsive transcriptional regulator (Wang et al., 2022), as well as global transcriptional regulators, including RsmA, GacA, RpoN1, and RpoN2, was modulated in NGJ1 strains. In *Burkholderia thailandensis,* OxyR regulates the secretion of T6SS effectors to scavenge manganese ions, thereby providing protection against oxidative stress (Si et al., 2017). GacA/RsmA regulatory network is a master regulator of secondary metabolism, stress adaptation, and secretion systems (Dupont et al., 2025; Wei et al., 2013), while RpoN is a sigma factor, known to play a multifaceted role in bacteria (Gu et al., 2022; Shao et al., 2018; Sun et al., 2025). Given that RpoN is involved in regulating T6SS in *Pseudomonas aeruginosa* (Allsopp et al., 2022b), we focused on characterizing RpoN’s involvement in modulating T6SS and redox homeostasis in NGJ1. We observed that RpoN1 is upregulated in the Δ*thio* mutant, and that the Δ*rpoN1* mutant has a nonfunctional T6SS, whereas overexpression of RpoN1 (RpoN1^OE^) activates the T6SS, even in the absence of the prey. Moreover, Δ*rpoN1* mutant was hypersensitive to oxidative stress, while the RpoN1^OE^ strain was able to manage the stress. Overall, our data suggest that oxidative stress signaling is induced in Δ*thio* mutant, leading to induction of *rpoN1* expression, which in turn triggers T6SS. Recent studies have revealed that in addition to its role in interbacterial competition, T6SS also contributes to oxidative stress tolerance by secreting effectors for the acquisition of essential transition metals, such as Zn^2+^ and Mn^2+^, that act as cofactors for antioxidant enzymes involved in redox detoxification (Si et al., 2017; Zuo et al., 2023). Thus, activation of the T6SS might aid intracellular ROS detoxification in NGJ1, in addition to its traditional array of functions.

The constitutive activation of the T6SS following Thioesterase dysfunction appears to be an adaptive response that couples metabolic imbalance with enhanced antibacterial competition. We speculate that integration of metabolic and stress-responsive pathways enables bacterial cells to fine-tune their antagonistic behavior in response to their physiological state within the complex microbial community. Our study reveals an unexplored interplay among secondary metabolism, redox homeostasis, T6SS-mediated redox detoxification, and antibacterial competition, highlighting secondary metabolite clusters as potential regulators of bacterial adaptation within the host.

## Material and method

### Growth conditions

*Burkholderia gladioli* strain NGJ1 and its derivatives were grown in PDA (Potato dextrose media, Himedia India) plates at 28°C. The bacterial strains used in this study are listed in the (Supplementary Table 2). The insertional mutants and complement strains of NGJ1 were grown on PDA supplemented with antibiotics. *Escherichia coli* was grown on LBA (Luria-Bertani Agar; HiMedia, India) plates at 37°C. *Xanthomonas oryzae* pv. *oryzae* strain (Xoo; BXO43 strain) was grown on PSA (Peptone Sucrose Agar, Himedia India) plates at 28°C (Jha et al. 2007). *Rhizoctonia solani* AG1-IA (strain BRS1) was grown on PDA (Himedia) plates at 28°C (Ghosh et al., 2021). *Oryza sativa* L. PB1 were grown in pots under greenhouse conditions (28 °C temperature, 80% relative humidity, 16/8 hr day/night cycle, and 250 µmol light intensity (Tyagi et al., 2024).

### *In silico* prediction of Biosynthetic gene cluster (BGCs)

Using antiSMASH 5.0 (https://antismash.secondarymetabolites.org), a Biosynthetic Gene Cluster (BGC) was predicted using the available draft genome sequence of NGJ1 (Jha et al. 2015), available at the Burkholderia genome database (https://www.burkholderia.com). The software predicts secondary metabolite-encoding BGCs using manually curated and validated gene cluster profile Hidden Markov Models (pHMMs) trained on multiple sequence alignments of empirically characterized proteins or protein domains(Blin et al., 2017).

### Insertional mutagenesis through plasmid integration

Insertional mutants were generated by homologous recombination-mediated plasmid integration as described previously (Das et al., 2022). Briefly, a partial fragment of the target gene was amplified from the genomic DNA of *B. gladioli* NGJ1 using gene-specific primers and cloned into the suicide vector pCZ367. The resulting construct was introduced into electrocompetent wild-type NGJ1 cells by electroporation using a Gene Pulser Xcell™ system (Bio-Rad). Transformants were selected on PDA supplemented with rifampicin (50 μg/ml), ampicillin (50 μg/ml), and gentamicin (50 μg /ml). Gus positive cells were selected as described by (Ray et al., 2015). The plasmid integration and disruption of the target gene were verified by PCR using a gene-specific flanking primer and the vector-specific primer.

### Generation of thioesterase-complement and *RpoN1^OE^ strains*

The full-length *thioesterase* gene was PCR-amplified from NGJ1 genomic DNA using the primer pair provided in the (Supplementary table 3). The amplified product was cloned into broad-host-range pHM1 vector, and the recombinant plasmid (pHM1::thio) was electroporated into Δ*thio*, to obtain the complement strain (Δ*thio+pHM1*::*thio*). The complement strain was confirmed by colony PCR using full-length gene-specific primers.

For the generation of the *RpoN1^OE^* strain, full-length *rpoN1* was amplified using NGJ1 genomic DNA and cloned into the *pHM1* vector. The recombinant pHM1::*rpoN1* plasmid was electroporated into WT NGJ1. The positive colonies were selected on rifamycin and spectinomycin and further confirmed by colony-PCR using full-length gene-specific primers and vector-specific primers.

### Bacterial-fungal confrontation assay

Bacterial–fungal confrontation assays were performed as described previously (Kumar et al., 2018; Yadav et al., 2020). Briefly, sclerotia of *Rhizoctonia solani* AG1-IA strain BRS1 were incubated with 5 mL of bacterial suspension (10 CFU mL⁻¹) of different NGJ1 strains for 4 h at 28 °C. Subsequently, the sclerotia were washed thoroughly with sterile water, transferred to fresh potato dextrose agar (PDA) plates, incubated at 28 °C, and growth was recorded at 4 and 7 days post-inoculation (dpi). Sclerotia treated with potato dextrose broth (PDB) served as the negative control. To assess sclerotia viability, secondary sclerotia recovered from the confrontation plates were transferred to fresh PDA plates and incubated at 28 °C to evaluate their germination.

### Biocontrol assay

The biocontrol efficacy of different NGJ1 strains in preventing *R. solani* infection in rice was performed as described previously(Das et al., 2022) . Briefly, *R. solani* sclerotia were incubated with bacterial suspensions (10 CFU mL⁻¹) for 4 h at 28 °C, washed thoroughly with sterile water, and inserted between the sheaths of rice (*Oryza sativa* L. cv. PB1) tillers using sterile forceps. Disease symptoms were assessed at 5 dpi, and severity was quantified by measuring Relative vertical sheath colonization (RVSC) as described in (Sahoo et al., 2026). The experiment was repeated at least three times, and for each experiment, at least 15 rice sheaths were analyzed per treatment.

### Inter-bacterial competition assay

Interbacterial competition assays were performed as described previously (Yadav et al., 2021). Briefly, cultures of NGJ1 (or its derivatives) and the prey bacteria (*E. coli* and Xoo) were adjusted to equal cell densities and mixed at a 1:1 ratio. The mixtures were spotted onto LB agar plates for *E. coli* or PSA plates for Xoo and incubated at 28°C for 24 h. Following incubation, the spotted bacterial cells were resuspended in 1 ml of LB broth, serially diluted, and plated onto selective agar media and growth temperature appropriate for the prey bacteria to enumerate surviving cells. Colony-forming units (CFUs) of the prey bacteria recovered from co-cultures were compared with those of prey bacteria incubated in the absence of NGJ1 or its derivatives, which served as the control. The experiment was independently repeated using at least three biological and technical replicates and similar results were observed in each replicate.

### *Xanthomonas oryzae* pv. oryzae infection

NGJ1 (or its derivatives) and Xoo cultures were grown separately in PDB and PSB, respectively, supplemented with the appropriate antibiotics, until an O.D of 1.0 was reached. Bacterial cells were harvested by centrifugation at 6,000 × *g* for 10 min, washed with sterile phosphate-buffered saline (PBS), and resuspended in PBS to a final density of approximately 10^9^ cells/mL. Equal volumes of NGJ1 (or its derivatives) and Xoo suspensions were mixed immediately before inoculation. Healthy leaves of 40–45-day-old rice plants (*Oryza sativa* L. PB1) were inoculated using the leaf-clipping method with sterile scissors dipped in the bacterial suspensions (Jha et al., 2007). Lesion lengths were measured 14 days post-inoculation to assess disease severity. At least 15 leaves were inoculated for each treatment in each experiment, and all experiments were performed at least three times independently.

### Hcp-1 secretion assay for T6SS activation

Total proteins were isolated from overnight cultures of different NGJ1 strains grown either alone or co-cultured with the prey bacterium (*E. coli* DH5α) at a predator-to-prey ratio of 5:1, as described previously (Yadav et al., 2021). Following co-incubation, cultures were centrifuged at 12,000 × *g* to obtain cell-free supernatants. Secreted proteins were precipitated from the supernatants by the addition of trichloroacetic acid (TCA; 12% w/v), washed twice with ice-cold 70% ethanol, air-dried, and resuspended in 1 ml of 10 mM PBS. Equal amounts of protein (10 μg) were separated on 12% SDS–PAGE gels and transferred onto PVDF membranes as described previously (Swain et al., 2017). Membranes were probed with rabbit polyclonal anti-Hcp1 (T6SS-1) antibody (ABGENEX Pvt. Ltd., India) diluted 1:25,000, followed by alkaline phosphatase-conjugated anti-rabbit IgG secondary antibody (Sigma–Aldrich) diluted 1:10,000. Bands were visualized using the NBT/BCIP substrate (Thermo Scientific, Cat. No. 34042) according to the manufacturer’s instructions.

### RNA Isolation, cDNA synthesis, and expression analysis

Total RNA was isolated from different NGJ1 strains using Qiagen RNA Isolation Kit as per manufacturer’s instructions, and 1 μg was used for cDNA synthesis using verso cDNA synthesis kit (Thermo Fisher Scientific Inc., Waltham, MA, USA). The qRT-PCR was performed by (PowerUp SYBR Green master mix (Applied Biosystems), according to the manufacturer’s instructions on ABI 7900HT™ Real-time PCR (Applied Biosystems). The gene-specific primers were used for expression analysis, as listed (Supplementary Table S3). The 16S rRNA gene of NGJ1 was used as a reference gene to normalize gene expression. The fold change in gene expression during the bacterial-fungal interaction at 24h was estimated using the 2^-ΔΔCt method (Livak & Schmittgen, 2001). The relative expression was calculated by 2^−ΔCt^. The experiments were independently repeated three times, using a minimum of three technical replicates.

### H_2_O_2_ Sensitivity Assay

Sensitivity to oxidative stress was assessed using a hydrogen peroxide (H₂O₂) disc diffusion assay as described previously (Pedre et al., 2018; Yu et al., 2016). Overnight cultures of the different NGJ1 strains were grown to an O.D. of 0.6 and uniformly spread onto PDA plates. Sterile paper discs were placed at the center of each plate and loaded with H₂O₂ solutions at the indicated concentrations (5mM or 20mM). Plates were incubated at 28°C for 48 h, and H₂O₂ sensitivity was determined by measuring the diameter of the inhibition zone. Further, different NGJ1 strains were grown in liquid medium with or without 20 mM H₂O₂. Following incubation at 28°C, bacterial growth was quantified by measuring optical density at 595 nm. Each experiment was independently repeated at least three times, with three technical replicates per biological replicate.

### Qualitative analysis of intracellular Reactive Oxygen Species (ROS) by Fluorescence spectroscopy and Confocal microscopy

Intracellular reactive oxygen species (ROS) accumulation was assessed using the ROS-sensitive fluorescent probe 2′,7′-dichlorodihydrofluorescein diacetate (H₂DCFDA) as described previously (Ghosh et al., 2017; Kim & Xue, 2020). The different NGJ1 strains were cultured in the presence or absence of H₂O₂ (20 mM) for 24 h. Cells were harvested by centrifugation at 5,000 × *g* for 5 min, resuspended in PBS (10 mM; pH 7.4), and incubated with 10 μM H₂DCFDA (Thermo Fisher Scientific Inc. USA, USA) for 30 min at room temperature in the dark. Following staining, cells were washed twice with PBS to remove excess dye. Intracellular ROS levels were quantified by measuring the fluorescence intensity of dichlorofluorescein using a fluorescence spectrophotometer (excitation, 485 nm; emission, 535 nm). Stained cells were also visualized using a Leica TCS SP8 AOBS confocal laser scanning microscope (Leica Microsystems, Wetzlar, Germany) under the GFP channel. Images were acquired and analyzed using LAS AF software (version 2.6.0, build 7266; Leica Microsystems). All experiments were performed using three independent biological replicates, each with three technical replicates.

## Statistical Significance

Statistical tests were performed using GraphPad Prism 10.0. Error bars indicate mean ±SE. Multiple means were compared using one-way ANOVA and Tukey’s multiple comparisons post hoc test, Two-way ANOVA and Tukey’s multiple comparisons post hoc test. Different letters indicate significant difference (p<0.05). Asterisks ^∗, ∗∗, ∗∗∗, ∗∗∗∗^ indicating a significant difference at p<0.5, p<0.05, p<0.001, and p<0.0001 respectively; were mentioned in the figure legend, wherever required.

## Acknowledgement

R.Y and M.R acknowledge fellowship from CSIR, Govt of India. G.J acknowledges funding from DBT under research projects: BT/PR4429/AGIII/103/1349/2021; BT/PR61470/CRA/168/1/2025 and RAD-42/15/2025-PBB-DBT and SR-NBACD award: HRD-20/3/2024-HRD-DBT. GJ also acknowledges Swarna Jayanti Fellowship (SB/SJF/2020-21/01) from ANRF (SERB). The authors are also thankful to DBT-eLibrary Consortium (DeLCON) for providing access to e-resources.

## Author Contribution

GJ conceived the work, supervised and coordinated its progress. RY initiated the work and identified BGCs clusters. KK helped in mutant characterization. RY and KK established the role of thioesterase in T6SS-medaited antibacterial activity. RK and RY performed confocal microscopy and western blot analysis. MR and RY performed qRT-PCR analysis. GJ, RY, and KK have contributed to writing the manuscript, and all authors have approved the final manuscript.

## Data Availability

The authors declare that all relevant data supporting the findings of the study are available in this article and its Supporting Information files.

## Supplementary information

**Figure S1:** Schematic representation of biosynthetic gene clusters(BGCs) in *B. gladioli* strain NGJ1.

**Figure S2:** The expression profile of selected genes of different BGCs during mycophagous interactions of *B. gladioli* strain NGJ1 with *R. solani*

**Figure S3:** Disruption of selected genes in BCGs compromises mycophagous ability of *B. gladioli* strain NGJ1.

**Figure S4:** Antibacterial activity of different *B. gladioli* NGJ1 strains.

**Figure S5.** Disruption of *thioesterase* gene compromises mycophagous and biocontrol ability of *B. gladioli* strain NGJ1.

**Figure S6:** Chrome azurol S (CAS) blue agar plate for siderphore estimation in NGJ1, Δthio and complemented strains after incubation at 28° C for 48h.

**Figure S7:** Sensitivity of different strains of NGJ1 against oxidative stress.

**Figure S8:** Sensitivity of Δ*mfs* mutant against oxidative stress.

**Supplementary Table 1:** The Biosynthetic Gene Cluster (BGCs) in B. gladioli strain NGJ1

**Supplementary Table S2:** Bacterial strains, plasmids, and antibodies used in this study

**Supplementary Table S3:** Primer list

