## Supplementary Table and Figure for "Thioesterase modulates antibacterial Type VI secretion system via alteration in oxidative stress responses in *Burkholderia*"

**Supplementary Table S1**: The Biosynthetic Gene Cluster (BGCs) in *B. gladioli* strain NGJ1

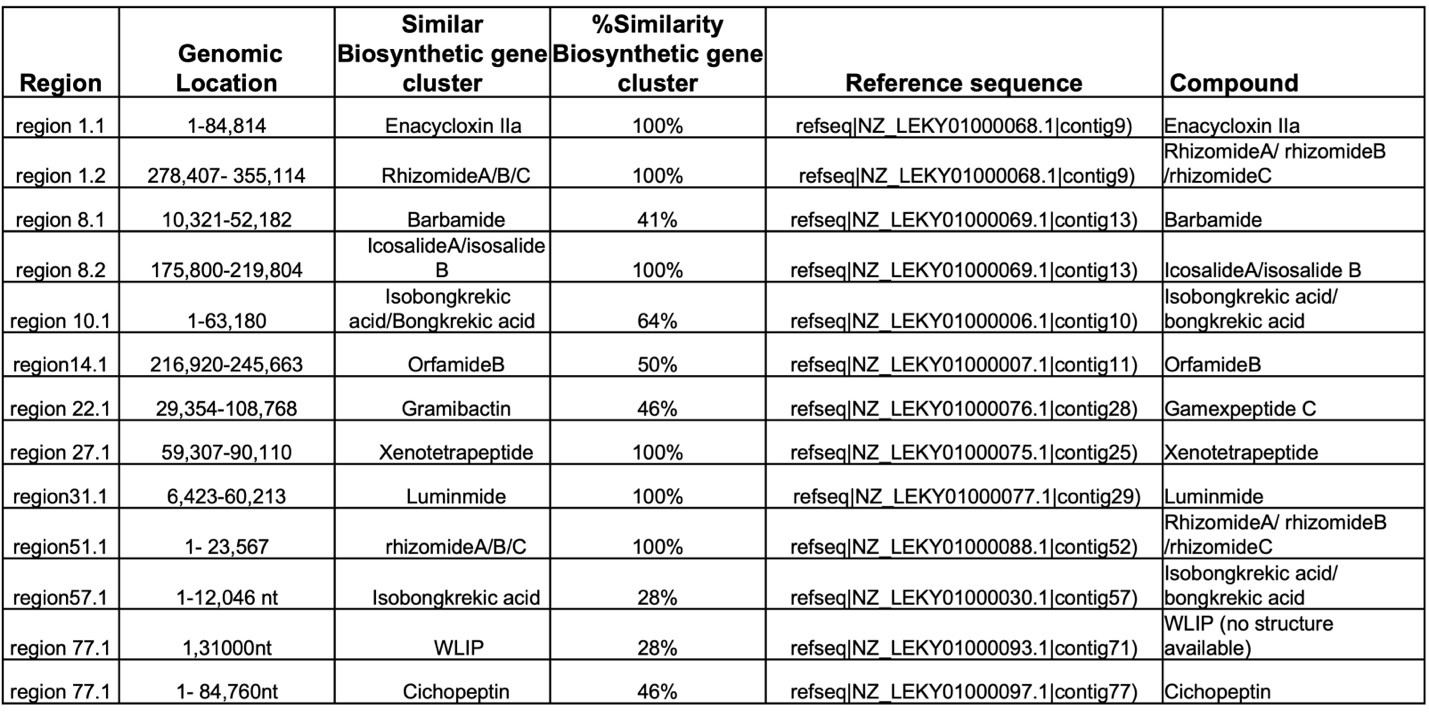

**Supplementary Table S2**: **Bacterial strains, plasmids, and antibodies used in this study**

| **Strains or plasmid** | **Relevant characteristics/ Source** | **Reference** |
| --- | --- | --- |
| ***E. coli* strains** |  |  |
| DH5α | F', endA1 hsdR17 (rk- mk+) supE44 thi-1 recA1 gyrA TelA1 cp8OdlacZAM15 A (lacZY A-argF)  U169 | Lab collection |
| BL21(DE3) | F- ompT gal dcm lon hsdSB(rB- mB-) λ(DE3 [lacI lacUV5-T7 gene 1 ind1 sam7 nin5]) | Lab collection |
| **Xanthomonas strain** |  |  |
| *Xanthomonas oryzae* BXO43 | Rif^r^ derivative | Lab Collection |
| ***B. gladioli* strains** |  |  |
| NGJ1 | Rif^r^ derivative of *B. gladioli* strain NGJ1 | Lab Collection  (Swain et al. 2017) |
| *Δthioesterase* | pCZ367::thio; Am^r^, Gm^r^ derivative of NGJ1 | Current Study |
| *Δmfs* | pCZ367::MFS-Transporter; Am^r^, Gm^r^ derivative of  NGJ1 | Current Study |
| *Δlux R* | pCZ367::Lux R ; Am^r^, Gm^r^ derivative of NGJ1 | Current Study |
| *Δmfet* | pCZ367::Mate Transporter; Am^r^, Gm^r^ derivative of  NGJ1 | Current Study |
| *Δcyp450* | pCZ367::CPY P450 ; Am^r^, Gm^r^ derivative of NGJ1 | Current Study |
| *Δpkb-acps* | pCZ367::ACP ; Am^r^, Gm^r^ derivative of NGJ1 | Current Study |
| NGJ201 (T6SS-1-) | *TssB*::pK18mob, Kmr derivative of NGJ-2 (T6SS-1 mutant strain) | Lab Collection  (Yadav et al. 2021) |
| Δ*rpoN1* | pK18mob::*rpoN1*; Km^r^ derivative of NGJ1 | Lab Collection  (Das et al. 2022) |
| *Δthio/pHM1::thio* | *Δthio/pHM1::thio;* Spr derivative of Δthio mutant strain  (Complement Strain) | Current Study |
| *ΔNGJ1/pHM1::rpoN1* | *NGJ1/pHM1::rpoN1;* Spr derivative of Δthio mutant strain (RpoN1^0.E^) | Current study |
| **Plasmid** |  |  |
| pCZ367 | Insertional vector with lacZ reporter (7.031kb); Am^r^,  Gm | Lab collection |
| pHM1 | Broad-host-range cosmid vector (13.3 kb); Sp^r^ | Lab collection |
| pET28a | Expression vector Kan^r^, Expression vector | Lab Collection |
| pCZ367::Thioesterase | Amp^r^, Gm, pCZ367 harbouring partial fragment of  Thioesterase | Current Study |

| pCZ367::MFS-Transporter | Amp^r^, Gm, pCZ367 harbouring partial fragment of  MFS-Transporter | Current Study |
| --- | --- | --- |
| pCZ367::CPY P450 | Amp^r^, Gm, pCZ367 harbouring partial fragment of CPY  P450 | Current Study |
| pCZ367::PKB-ACP | Amp^r^, Gm, pCZ367 harbouring partial fragment of ACP | Current Study |
| pCZ367::Lux R | Amp^r^, Gm, pCZ367 harbouring partial fragment of Lux  R | Current Study |
| pCZ367::Mate Transporter | Amp^r^, Gm, pCZ367 harbouring partial fragment of  Mate Transporter | Current Study |
| pHM1:RpoN1 | Sp^r^ ,pHM1 harbouring full fragment of RpoN1 | Lab collection (Das et al.,2022) |
| pHM1:Thioesterase | Sp^r^ ,pHM1 harbouring full fragment of Thioesterase | Current Study |
| pET28a::Thioesterase | Kan^r^, pET28a harbouring full fragment of Thioesterase | Current Study |
| **Antibodies** |  |  |
| Polyclonal peptide anti-  Hcp-1 raised in Rabbit | ABGENEX Pvt. Ltd., India | Current study |
| Anti-Rabbit IgG tagged with alkaline phosphatase | (Sigma, Catalog no.- A3812) | Current study |

**Supplementary Table S3: Primer list**

| **Gene** | **Primer Sequence** | **Reference** |
| --- | --- | --- |
| **For Insertional Mutant creation** | |  |
| MFET partial fragment cloning in pCZ-367 | Forward: TCTAGACTTCCGCATCTACGTCTATT | Current Study |
|  | Reverse: AAGCTTACGATCAGCGAATAGATCAC | Current Study |
| Lux R partial fragment cloning in pCZ-367 | Forward: TCTAGACCTATTACCTCCATACGCCC | Current Study |
|  | Reverse: AAGCTTCGTTCGAGGTATTGAGCTTG | Current Study |
| CYP450 partial fragment cloning in pCZ-367 | Forward: TCTAGACAATACCTGGCTGATGTTCC | Current Study |
|  | Reverse: AAGCTTATTCCTCCTCGCTCATCTG | Current Study |
| PBK-ACPs partial fragment cloning in pCZ-367 | Forward: TCTAGACGCGCCAATTCGGTATCAAG | Current Study |
|  | Reverse: AAGCTTGAATGGATGCGGGTAAGGCA | Current Study |
| MFS-Transporter partial fragment cloning in pCZ-367 | Forward: TCTAGACTGGCACATCTACACGCTGA | Current Study |
|  | Reverse: AAGCTTGATGAAGGCGATGCCAGAGA | Current Study |
| Thioesterase partial fragment cloning in pCZ-367 | Forward: TCTAGAGCTGATCGCGCTGGA | Current Study |
|  | Reverse: AAGCTTCTTCGAGATAGAAATGGCCG | Current Study |
| Thioesterase full-length cloning in pHM1 | Forward: AAGCTTATGTCGGTCCACGCCGGC | Current Study |
|  | Reverse: GGTACCTCATTGCGTCCATCCTGCC | Current Study |
| RpoN1 full-length cloning on pHM1 | Forward: AAGCTTATGAAAGCCAGCCTCCAACT | Lab Collection |
|  | Reverse: GGTACCCTACAGAGACTTGCGCAGATTCA | Lab Collection |
| Thioesterase full-length cloned in pET-28 | Forward: GGATCCATGTCGGTCCACGCCGG | Current Study |
|  | Reverse: AAGCTTTTGCGTCCATCCTGCCTCC | Current Study |
| oLac Z | Forward: AAGGGGGATGTGCTGCAAGG | Current Study |
|  | Reverse: CAAAGCGCCATTCCCCATTC | Current study |
| **For qRT -PCR analysis** | |  |
| PKB-acp synthase | Forward: CTGATGGACCTGTCGCACTG | Current study |
|  | Reverse: TGAAACCGTCGTGATCGAGG | Current study |
| CPY P450 | Forward: CTGTCACACCTGTCGTTCCT | Current study |
|  | Reverse: CAGCCAGGTATTGGGCATCA | Current study |
| Lux R | Forward: TCCGCTGTCGATGAAGATCC | Current study |
|  | Reverse: AGACCAGATGGCACTGGAAG | Current study |
| Mate Transporter | Forward: ATCTGCTGCTGCTGGTGATT | Current study |
|  | Reverse: TCAGCGAATAGATCACGCCC | Current study |
| Thioesterae | Forward: TGGATCGACGAGGTCTG | Current study |
|  | Reverse: GCGAAGGCGTGGAAG | Current study |
| MFS-Transporter | Forward: GCTCAAGCTGAGCAGCAAAC | Current study |
|  | Reverse: GAAGGCGATGCCAGAGAACA | Current study |
| ACP | Forward: ATCGACTCGCTGAACATCGT | Current study |
|  | Reverse: TGCGTGAGGTTGATCTGCTC | Current study |
| Hyl-D | Forward: TCCTACGTACTCCCCGCTC | Current study |
|  | Reverse: TGATGGTCTTGGACAGCACG | Current study |
| Lys R | Forward: ACCTTCGCCATCAAGGAAA | Current study |
|  | Reverse: GTAGGGCCCGATCGTATAG | Current study |
| Gac A | Forward: TCAAGGACGAGAAACCGAGC | Current study |
|  | Reverse: GCTCTCCTTGGTCAGGTAGC | Current study |
| Rsm A | Forward: CACCTGATGAACTTCGCC | Current study |
|  | Reverse: GATCAGCTTGTCCATCACG | Current study |
| TseTBg1 | Forward: TTTCGAGATCCGCTGCATGA | Current study |
|  | Reverse: GGTCAGGAACCATTGAGCGA | Current study |
| 9Tle | Forward: GAATCAGGTTTGCCCTTGGC | Current study |
|  | Reverse: CACGCGAGAACCCGAAAATC | Current study |
| 12Tpe | Forward: AGTGTATGTTGCGCTCTCCC | Current study |
|  | Reverse: AGCCGCCGATTTGAAGTACA | Current study |
| 19Tde | Forward: TACCGTCCAGAAGTCGCCTA | Current study |
|  | Reverse: CCCGGCCAACTTCTTGTACT | Current study |
| 31Tae | Forward: GGTGATATCGAGCCTGACCG | Current study |
|  | Reverse: TAGTACGGCAAGCCTTCCAC | Current study |
| TssB | Forward: CCGCAAGTTCGTCAACATCG | Current study |
|  | Reverse: GAGCGAACGGAACTTGAGGT | Current study |
| TssJ | Forward: CGCCACGCAATTTAGGCATT | Current study |
|  | Reverse: GGGTCCTTCAGCGTGTAGAG | Current study |
| Hcp | Forward: AATCGCAGGACAAGGACCAC | Current study |
|  | Reverse: GTGAACACCATGTCGCTGTG | Current study |
| RpoN1 | Forward: CTTCGATCTCGAGCCTTCGG | Lab Collection |
|  | Reverse: ATTGAGCGGGTACAGGCAAA | Lab Collection |
| RpoN2 | Forward: ATTGTTGCGACTCGATCCGT | Lab Collection |
|  | Reverse: GCAATTCGAGATGCTGCGAG | Lab Collection |
| 16S rRNA | Forward: GGAGTACGGTCGCAAGATTAAA | Lab Collection |
|  | Reverse: GACCATGTCAAGGGTAGGTAAG | Lab Collection |

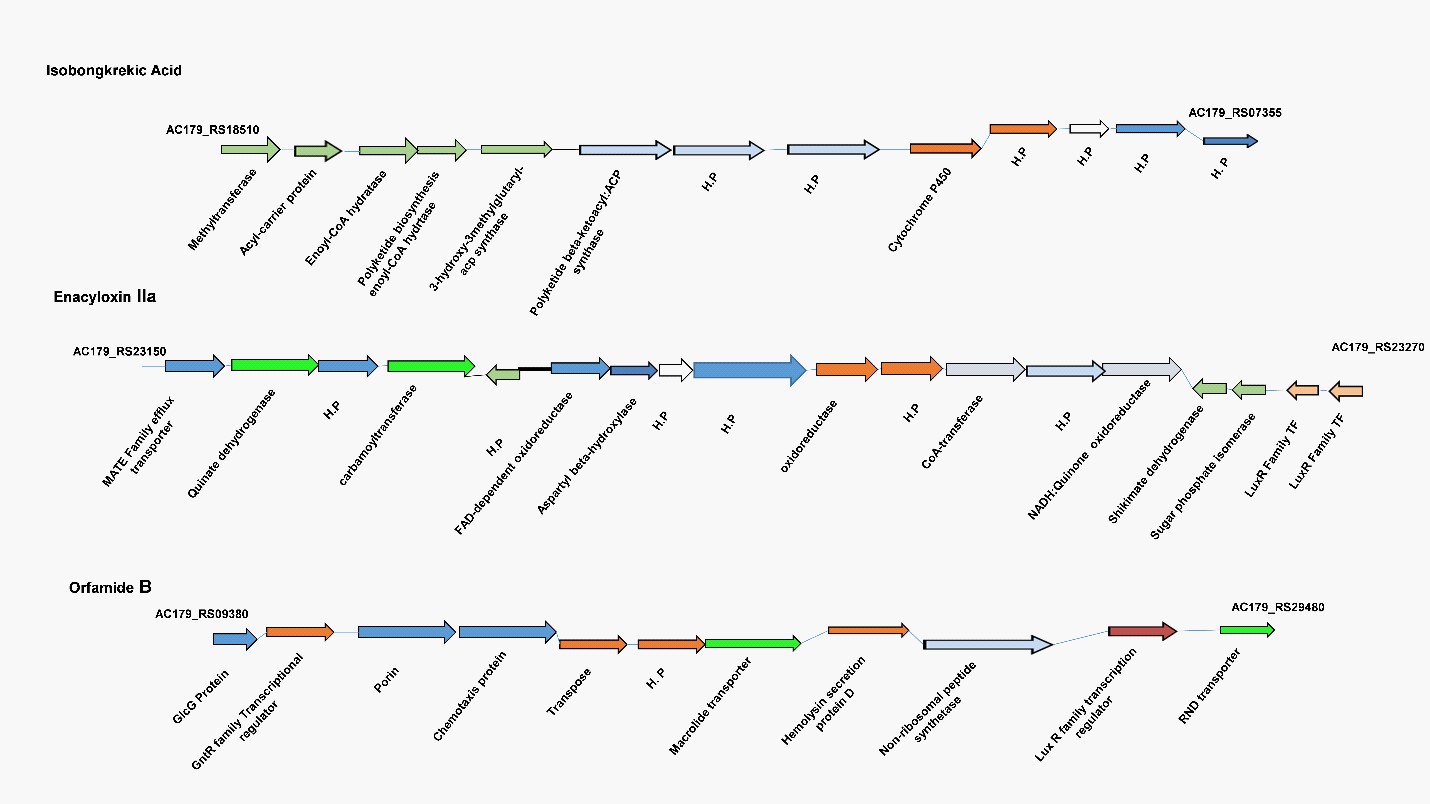

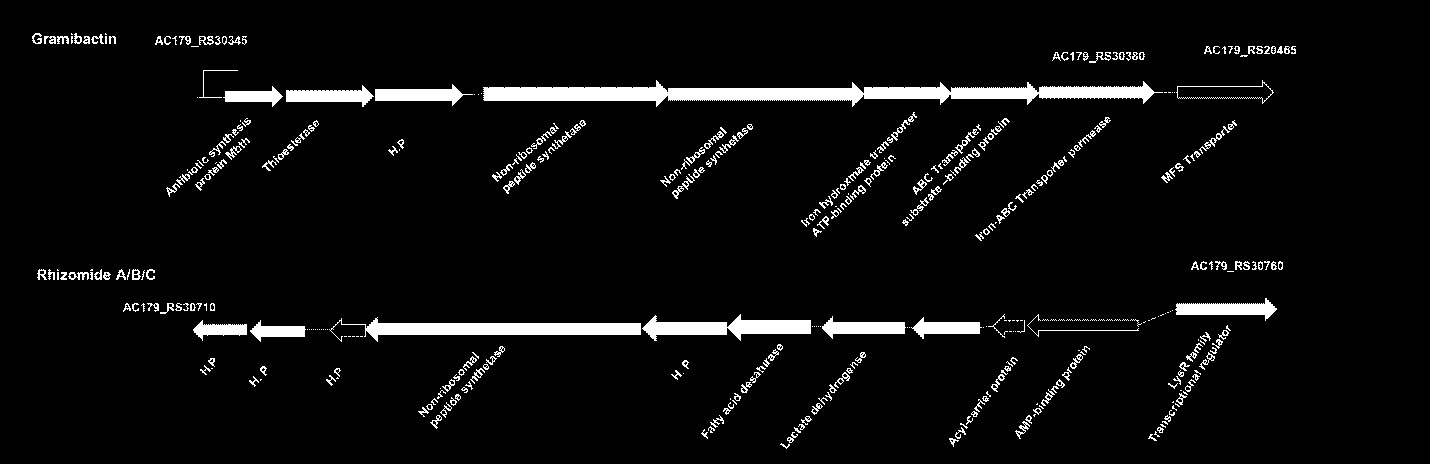

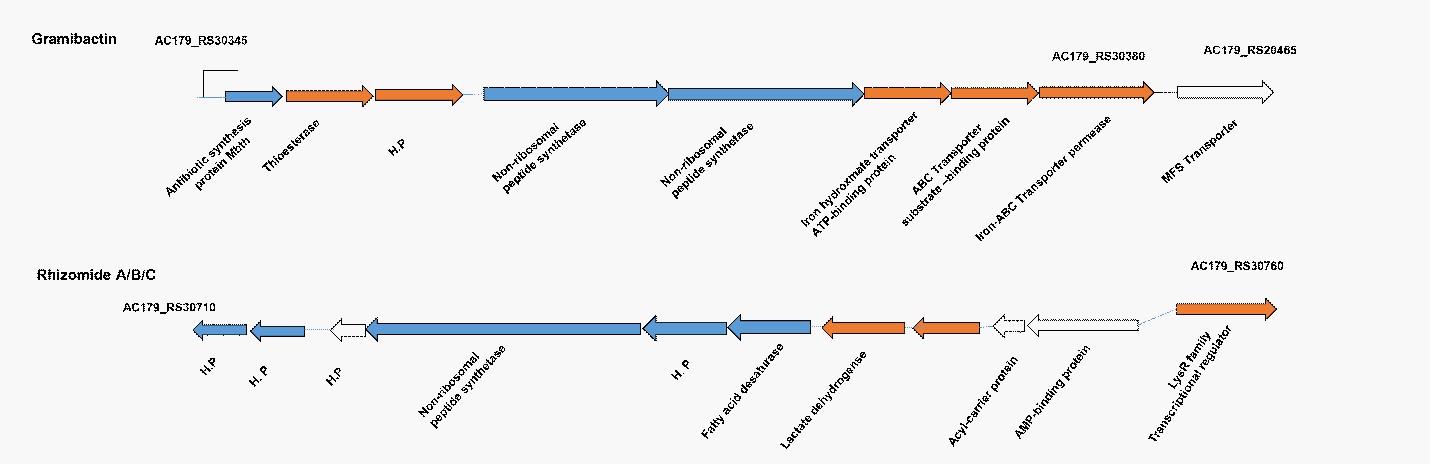

**Supplementary Figure S1.** Schematic representation of Isobongkrekic acid, Enacycloxin IIa, Orfamide B, Rhizomide A/B/C, encoding biosynthetic gene clusters in *B. gladioli* strain NGJ1.

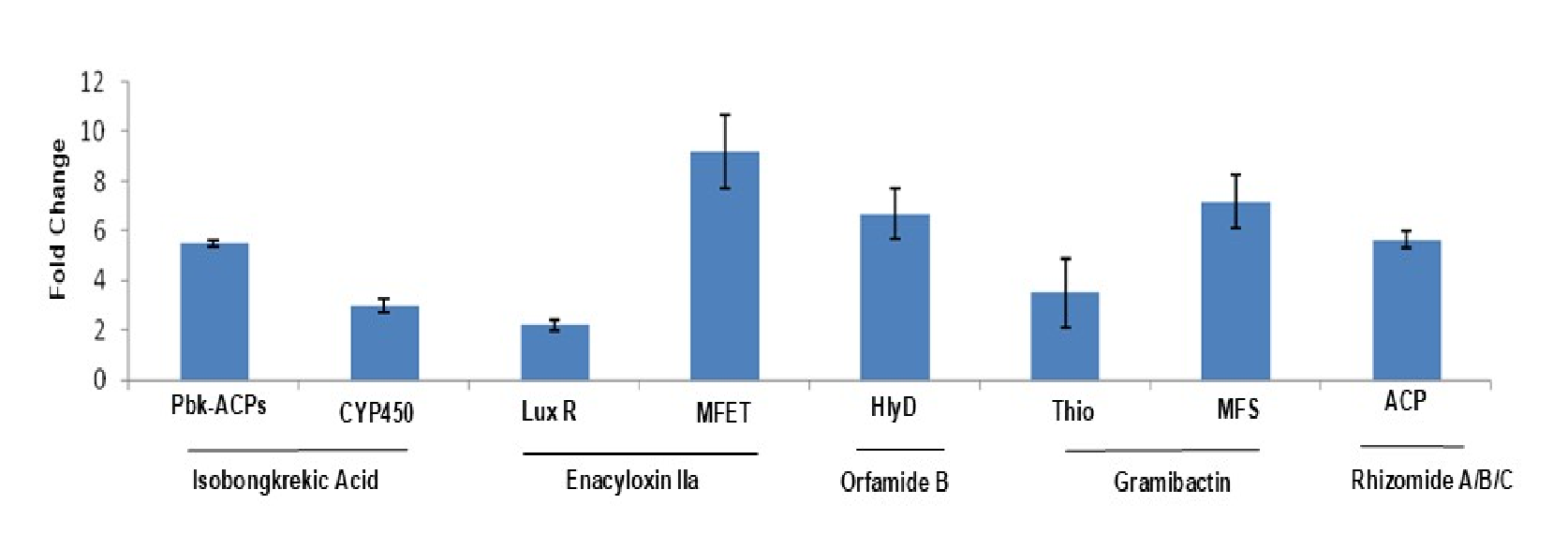

**Supplementary Figure S2**. **The expression profile of selected genes of different BGCs during mycophagous interactions of *B. gladioli* strain NGJ1 with *R. solani***. The qRT-PCR-based expression dynamics of genes encoding Pbk-ACPs synthase, and CYP450 of Isobongkrekic acid BGC, LuxR, and Mate Transporter of Enacyloxin IIa BGC, Hemolysin secretion protein D (HlyD) of Orfamide B BGC, Thioesterase and MFS transporter of Gramibactin BGC, and ACP gene of Rhizomide A/B/C BGC, during mycophagous interaction at 24h.The fold change in gene expression was calculated during mycophagous interactions with R. solani relative to gene expression during growth in laboratory media (without R. solani), using 16S rRNA from NGJ1 as an endogenous control. The experiments were independently repeated three times, with three technical replicates per repetition. The graph shows the mean ± standard error across three biological replicates.

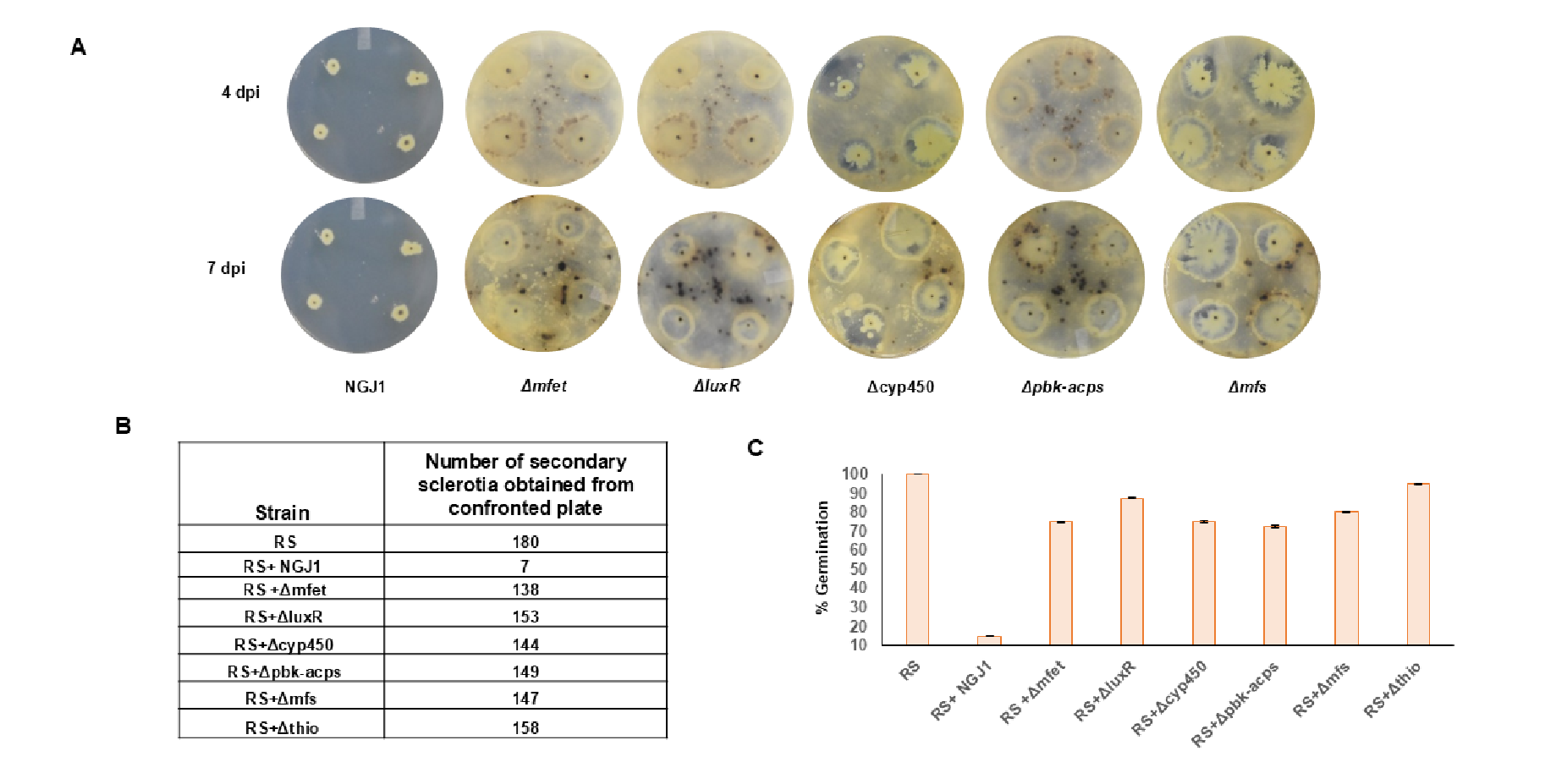

**Supplementary Figure S3. Disruption of selected genes in BCGs compromises mycophagous ability of *B. gladioli* strain NGJ1. A)** Representative images of mycophagous interaction of NGJ1 (WT) and mutant strains with *R. solani* on PDA plates, at 4 dpi and 7 dpi. **B)** Number of sclerotia produced by *R. solani* upon confrontation with NGJ1 strains on PDA plates, at 7 dpi**. C)** Germination rate (percentage) of sclerotia collected from bacterial-fungal confrontation plates, when transferred on fresh PDA plates. The experiments were independently repeated three times, with three technical replicates per repetition. The graph shows the mean ± standard error across three biological replicates.

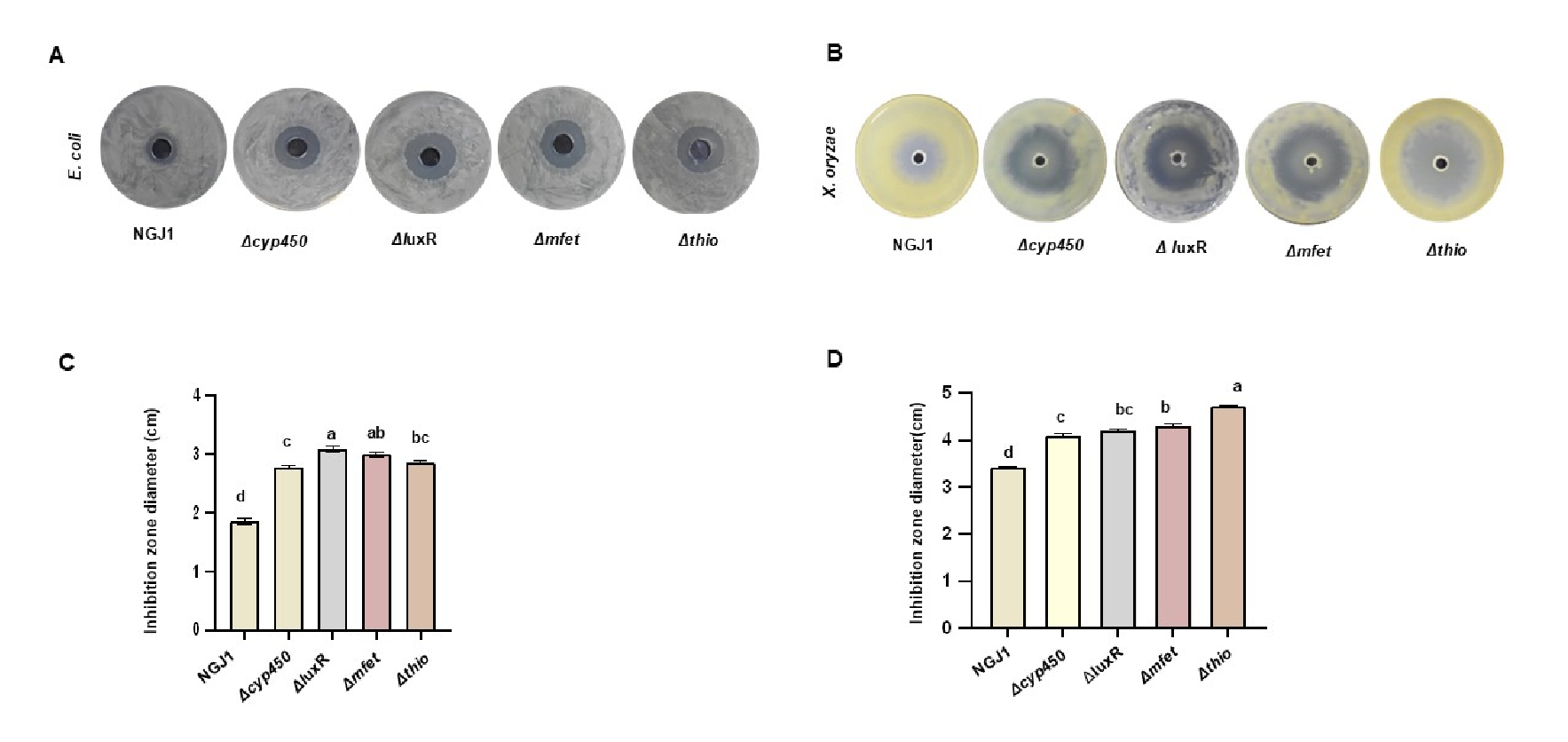

**Supplementary Figure S4. Antibacterial activity of different *B. gladioli* NGJ1 strains.** (**A**) and (**B**) represents the zone of inhibition caused by NGJ1 against *E. coli and X. oryzae,* respectively. (**C**) and (**D**) represents the diameter of the inhibition zone against *E. coli* and *X. oryzae,* respectively*.* The experiments were independently repeated three times, with a minimum of 3 technical replicates per condition. Data were analyzed using one-way ANOVA and Tukey's multiple comparisons post hoc test. Different letters represent significant differences (p<0.05). The graph shows the mean ± standard error across three biological replicates.

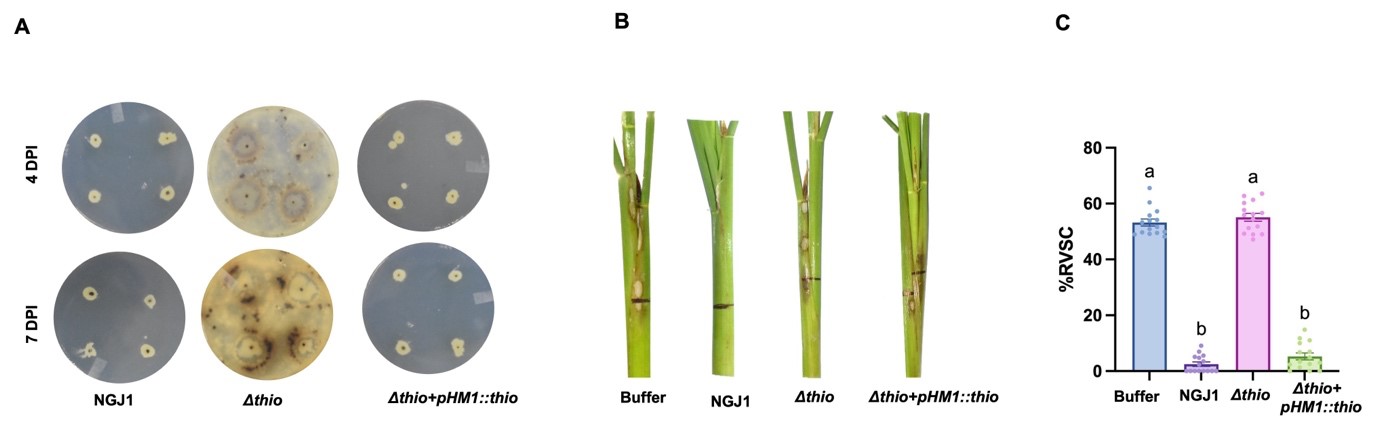

**Supplementary Figure S5. Disruption of *thioesterase* gene compromises mycophagous and biocontrol ability of *B. gladioli* strain NGJ1**. **A)** Representative photographs of mycophagous interaction of NGJ1 (WT), *Δthio* mutant, and complement (*Δthio+pHM1::thio*) strains with *R. solani* on PDA plates, at 4 dpi and 7 dpi. **B)** The biocontrol ability of WT, *Δthio*, and complement (*Δthio+pHM1::thio*) NGJ1 strains against *R. solani,* at 5 dpi*.* The sheath blight disease symptoms caused by infection with the bacterial-treated *R. solani* are shown. **C)** The Relative vertical sheath colonization (RVSC)-based disease index in rice. The Bar graph was analyzed by one-way ANOVA, Tukey’s multiple comparisons test, and post hoc test. Different letters represent significant differences (p<0.05). The experiments were independently repeated three times, with three technical replicates per repetition. The graph shows the mean ± standard error across three biological replicates.

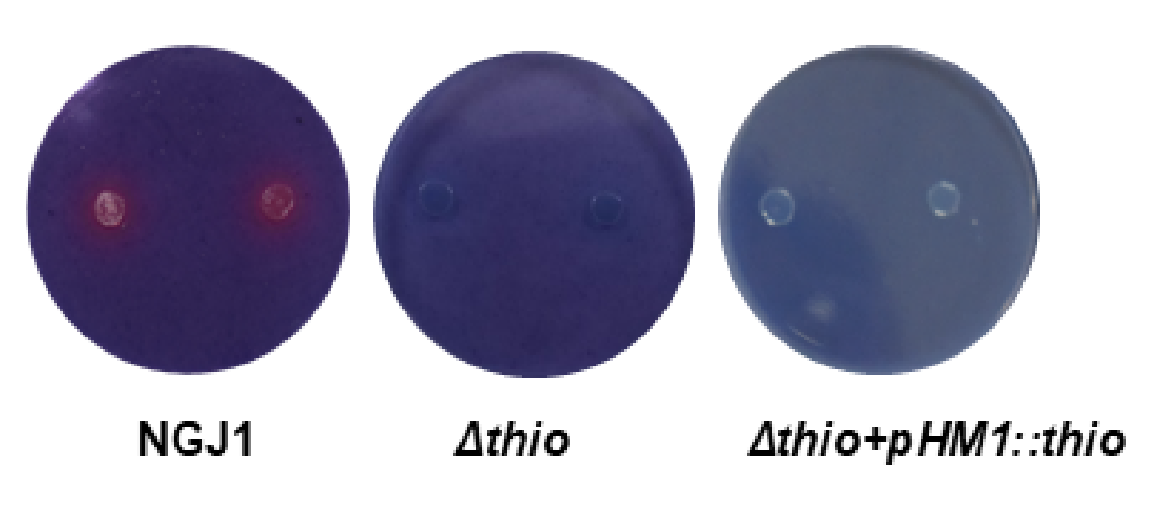

**Supplementary Figure S6.** Chrome azurol S (CAS) blue agar plate assay showing the production of siderophore by NGJ1, ∆thio mutant, and complemented strains.

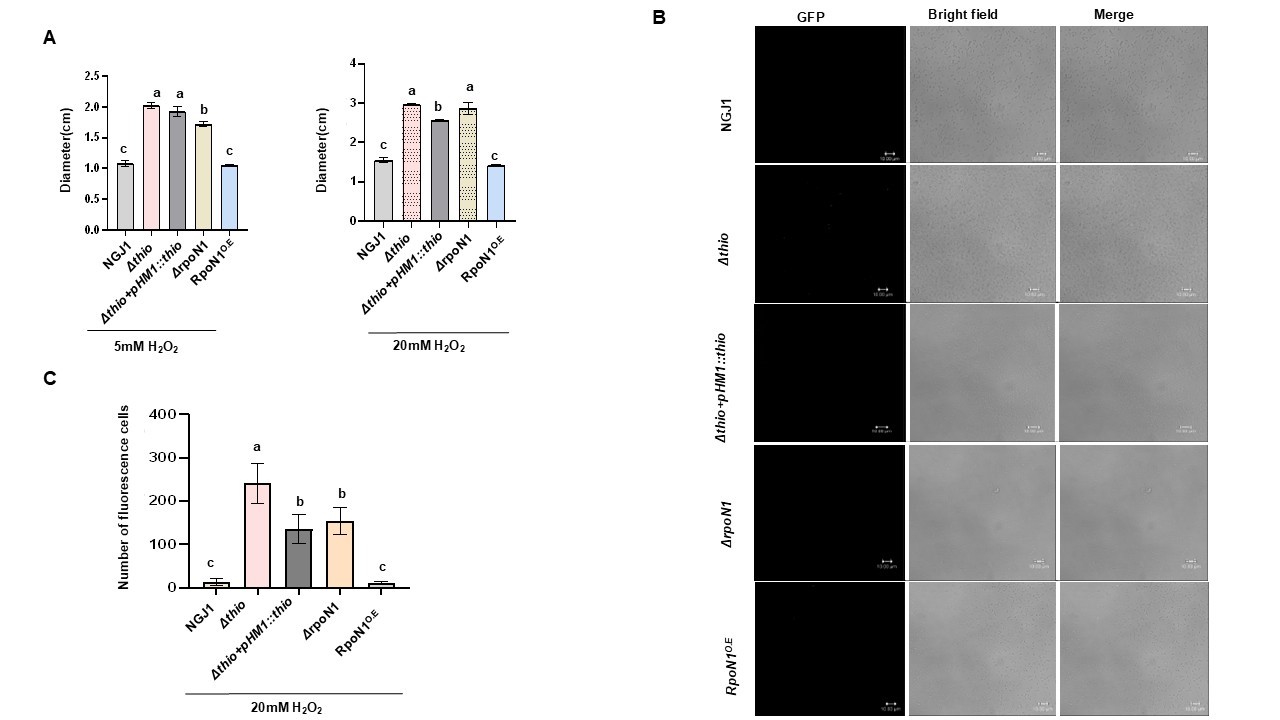

**Supplementary Fig 7. Sensitivity of different strains of NGJ1 against oxidative stress. A)** Graph showing the diameter of the inhibition zone in different strains of NGJ1 against 5 and 20mM H_2_O_2_ treatment_._ **B)** ROS accumulation (fluorescence signal, analyzed after H_2_DCFDA staining under a confocal microscope) in different strains of NGJ1, without H_2_O_2_ treatment. Scale bars = 10 µm. Similar results were obtained in at least 3 independent replicates, and only representative images are shown. **C)** The number of H_2_DCFDA-stained fluorescence cells observed in different NGJ1 strains upon

H_2_O_2_	treatment_._	Data	was	analysed	by	one-way	ANOVA,

Tukey's multiple comparisons test, and post hoc test. Different letters represent a significant difference (p<0.05). The graph shows the mean ± standard error across three biological replicates. The experiments were independently repeated three times, with a minimum of 3 technical replicates per condition. The graph shows the mean ± standard error across three biological replicates.

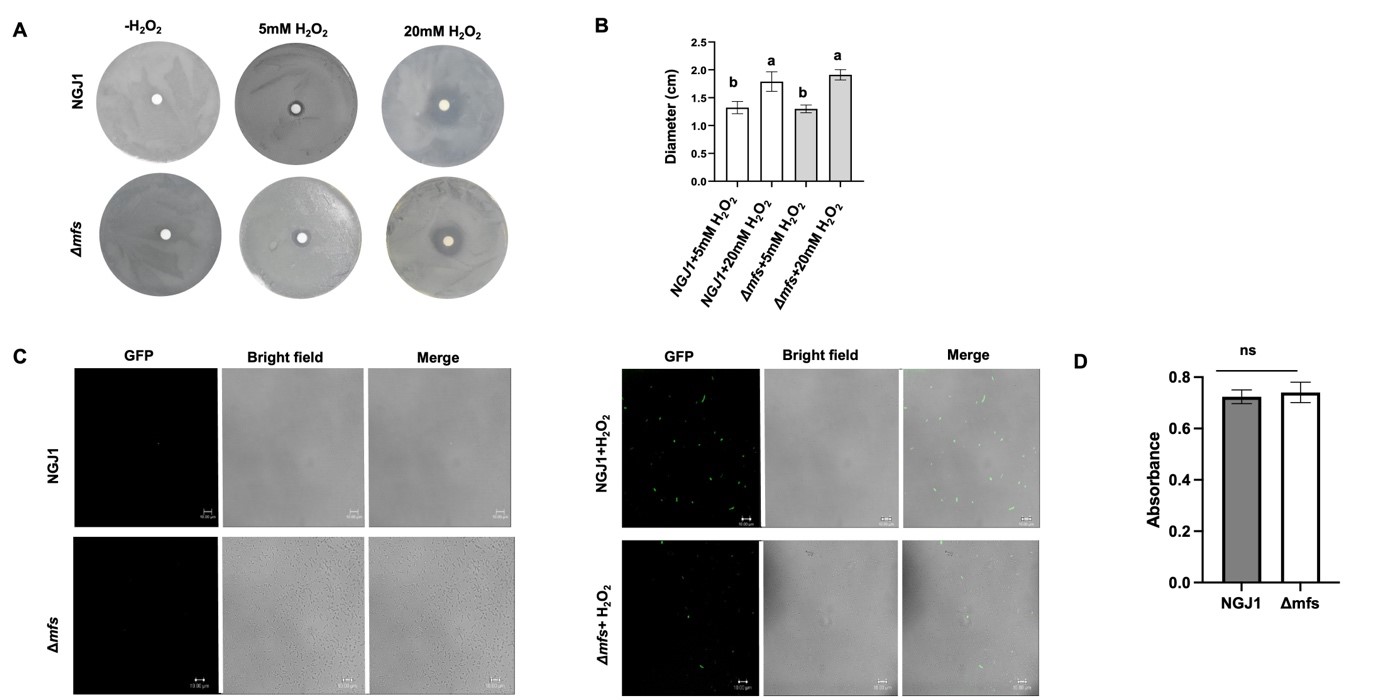

**Supplementary Fig S8. Sensitivity of *Δmfs* mutant against oxidative stress**. **A)** Sensitivity of NGJ1 (WT) and *∆mfs* strains, when grown for 48h on PDA plates supplemented with 5 and 20 mM H_2_O_2_. **(B)** Graph showing diameters of the inhibition zones of different NGJ1 strains. **(C)** Confocal micrograph showing ROS accumulation (Fluorescent cells upon H_2_DCFDA staining) in different strains of NGJ1, with and without 20mM H_2_O_2_ treatment, at 24h, scale bars = 10 µm. Similar results were obtained in at least 3 independent replicates, and only representative images are shown. **D)** The cell density of NGJ1 cells upon 24h of H_2_O_2_ treatment, measured as O.D^595nm^_._ Data was analyzed by one-way ANOVA and Tukey’s multiple comparisons post hoc test. Different letters represent significant differences (p<0.05). The graph shows the mean ± standard error across three biological replicates. The experiments were independently repeated three times with a minimum of three technical replicates
